# Phase Transition of RNA-protein Complexes into Ordered Hollow Condensates

**DOI:** 10.1101/2020.01.10.902353

**Authors:** Ibraheem Alshareedah, Mahdi Muhammad Moosa, Muralikrishna Raju, Davit Potoyan, Priya R. Banerjee

**Affiliations:** Department of Physics, University at Buffalo, Buffalo NY 14260, USA; Department of Chemistry, Iowa State University, Ames IA 50011, USA

**Keywords:** Nucleoprotein assembly, RNA vesicles, Optical tweezer, MD simulation, Biomolecular condensates

## Abstract

Liquid-liquid phase separation of multivalent intrinsically disordered protein-RNA complexes is ubiquitous in both natural and biomimetic systems. So far, isotropic liquid droplets are the most commonly observed topology of RNA-protein condensates in experiments and simulations. Here, by systematically studying the phase behavior of RNA-protein complexes across varied mixture compositions, we report a hollow vesicle-like condensate phase of nucleoprotein assemblies that is distinct from RNA-protein droplets. We show that these vesicular condensates are stable at specific mixture compositions and concentration regimes within the phase diagram and are formed through the phase separation of anisotropic protein-RNA complexes. Similar to membranes composed of amphiphilic lipids, these nucleoprotein-RNA vesicular membranes exhibit local ordering, size-dependent permeability, and selective encapsulation capacity without sacrificing their dynamic formation and dissolution in response to physicochemical stimuli. Our findings suggest that protein-RNA complexes can robustly create lipid-free vesicle-like enclosures by phase separation.

**Significance statement:** Vesicular assemblies play crucial roles in subcellular organization as well as in biotechnological applications. Classically, the ability to form such assemblies were primarily assigned to lipids and lipid-like amphiphilic molecules. Here, we show that disordered RNA-protein complexes can form vesicle-like ordered assemblies at disproportionate mixture compositions. We also show that the ability to form vesicular assemblies is generic to multi-component systems where phase separation is driven by heterotypic interactions. We speculate that such vesicular assemblies play crucial roles in the formation of dynamic multi-layered subcellular membrane-less organelles and can be utilized to fabricate novel stimuli-responsive microscale systems.

## Introduction

Precise spatiotemporal control of biochemical processes is indispensable to life. Classically, it was assumed that amphiphilic lipids provide living systems the capacity to segregate different biological processes into distinct membrane-bound compartments and therefore, afford spatial and temporal separation of (sub)cellular biochemistry (1). These membrane-bound compartments provide an independent internal environment that can be tuned as per cellular needs. Recent advances indicate that, in addition to membrane-bound organelles, cells also utilize membrane-free protein/RNA-rich condensates as compartments for organizing the intracellular space (2-5). These fluid compartments, often referred to as membrane-less organelles (MLOs), are formed by liquid-liquid phase separation (LLPS) and can partition a diverse set of biomolecules selectively (6). One key advantage of these MLOs is their dynamic nature wherein they can rapidly form and dissolve in a stimuli-responsive fashion.

Since many MLOs are thought to form by LLPS, their internal microenvironment is significantly higher in density as compared to the surrounding cytoplasm/nucleoplasm and exhibit complex fluid-like properties (7). To spatially segregate biochemical processes within this condensed phase, many MLOs [*e.g*., nuclei (8), nuclear speckles (9), paraspeckles (10), stress granules (11, 12) and P-granules (13)] utilize distinct sub-compartments within themselves. Such multilayered structures are best described by a coexisting multi-phasic condensate model where two or more distinct types of condensates are formed by LLPS of individual components in a multi-component mixture (5). However, a different class of multilayered MLOs has also been reported, where the layered topology is manifested due to the presence of a hollow internal space. For example, *in vivo* experiments have demonstrated that both nuclear and cytoplasmic germ granules of *Drosophilla* can exhibit hollow morphologies (14-16). In another system, it was observed that simple overexpression of TDP-43, a stress granule protein, can give rise to multilayered compartments with vacuolated nucleoplasm-filled internal space (17). However, physical driving forces behind these hollow morphologies remain poorly understood.

Recently, we demonstrated that RNA can mediate a reentrant phase transition of ribonucleoproteins containing arginine-rich low <u>c</u>omplexity <u>d</u>omains (LCDs) through multivalent heterotypic interactions (18, 19). At sub-stoichiometric regime, RNA triggers ribonucleoprotein (RNP) phase separation whereas at super-stoichiometric ratio, excess RNA leads to droplet dissolution due to charge inversion on the surface of RNP-RNA complexes (18). This inversion of charge suggests that the stoichiometry of these *fuzzy protein-RNA assemblies* is not fixed but varies with the mixture composition. The condensate dissolution at super stoichiometric ratio is expected to be generic for systems with obligate heterotypic interactions (20). Interestingly, we also observed that sudden jumps in RNA-to-LCD stoichiometry can lead to the formation of transient hollow droplet topologies under non-equilibrium conditions (18). Although, these vacuolated structures were observed only as fleeting intermediates [mean lifetime < 300 s] during reentrant dissolution of protein-RNA condensates (18), their occurrence nevertheless suggested that associative RNA-protein systems may have access to complex morphologies distinct from isotropic liquid droplets. This leads to the following question: are these hollow droplets only kinetic intermediates or are they representative of potentially stable topologies of two-component (excluding the solvent) associative condensates?

Here, to ascertain these possibilities, we explore the liquid-liquid phase separation regime within protein-RNA reentrant phase space that spans three orders of magnitude in concentrations. Using a combination of biophysical and computational tools (*e.g*., confocal microscopy, ensemble spectroscopy, optical tweezers, microfluidic manipulation, and molecular dynamics simulation), we report a hitherto unknown structural transition from isotropic liquid droplets to a vesicle-like phase in nucleoprotein-RNA condensates. These vesicle-like condensates are hollow structures enclosed by an ordered nucleoprotein-RNA membrane, and are formed through liquid-liquid phase separation of nucleoprotein-RNA complexes at distinct mixture compositions ([nucleotide]:[Arg] is approximately > 1.87 or < 0.075) and concentration regimes.

## Results and Discussion

### Nucleoprotein-RNA complexes form stable hollow condensates with vesicle-like properties

Our previous observations suggested that hollow condensate topology may exist when RNA concentration is substantially higher than RNP concentration (18). To investigate such complex morphologies, we first probed for the mesoscale structure of condensates formed by a model arginine-rich disordered nucleoprotein, protamine [PRM, see *SI Appendix*, Fig. S1*A*, (21)], with a single-stranded RNA [poly(U)] at excess RNA conditions (*C*_*RNA*_ *=* 5 *× C*_*PRM*_). Remarkably, we observed that micron-sized hollow condensates are readily formed by mixing 4.4 mg/ml PRM with 22 mg/ml RNA. These hollow condensates closely resemble lipid vesicles in their appearance (Fig 1*A*). Furthermore, we observed that these hollow condensates are relatively stable across a wide range of temperature and salt concentrations (*SI Appendix*, Fig. S2). Below 11 mg/ml RNA, PRM-RNA mixtures formed spherical liquid droplets that are uniform in density (Fig. 1*A* and *B, SI Appendix*, Fig. S3). Z-stack analysis using confocal fluorescence microscopy revealed that the hollow condensates have a rim and an internal lumen (Fig. 1*C, SI Appendix*, Fig. S4). PRM localizes within the rim but not in the lumen. We then asked whether RNA also localizes within the PRM-rich rim or inside the lumen. Two-color confocal microscopy revealed that RNA [probed by SYTO13 which fluoresces upon RNA binding (22)] co-localizes with PRM and is highly enriched in the rim. The lumen appears to be devoid of PRM and RNA relative to the rim as judged by the fluorescence intensity measurements radially through the hollow condensates (Fig 1*D, SI Appendix*, Fig. S5). To ascertain whether the lumen environment is indeed a *dilute phase* (as classical membrane-enclosed lumen environment is in lipid vesicles), we measured macromolecule diffusion (using TMR-labeled dextran as a probe) *outside, in the lumens*, and *within the rims* of hollow condensates utilizing fluorescence correlation spectroscopy (FCS; *SI appendix*, Fig. S6). FCS autocorrelation traces clearly show that probe diffusion is significantly slowed down within the rim (*e.g*., the diffusion half-time is increased by three orders of magnitude), whereas diffusion remains similar for molecules that are localized in the lumen as those in the external dilute phase (Fig. 1*E*). Similar FCS experiments with Alexa594-labeled PRM corroborated these results (*SI appendix*, Fig. S6A). Collectively, these experimental results indicate that (i) PRM-RNA complexes can form lipid-free vesicular structures at excess RNA-to-PRM ratio (*C*_*RNA*_ *> 2.5 × C*_*PRM*_), and (ii) the nucleoprotein membrane-enclosed internal space is a distinct *low-density phase* that is independent of the condensed fluid phase of vesicle rims.

**Figure 1:**
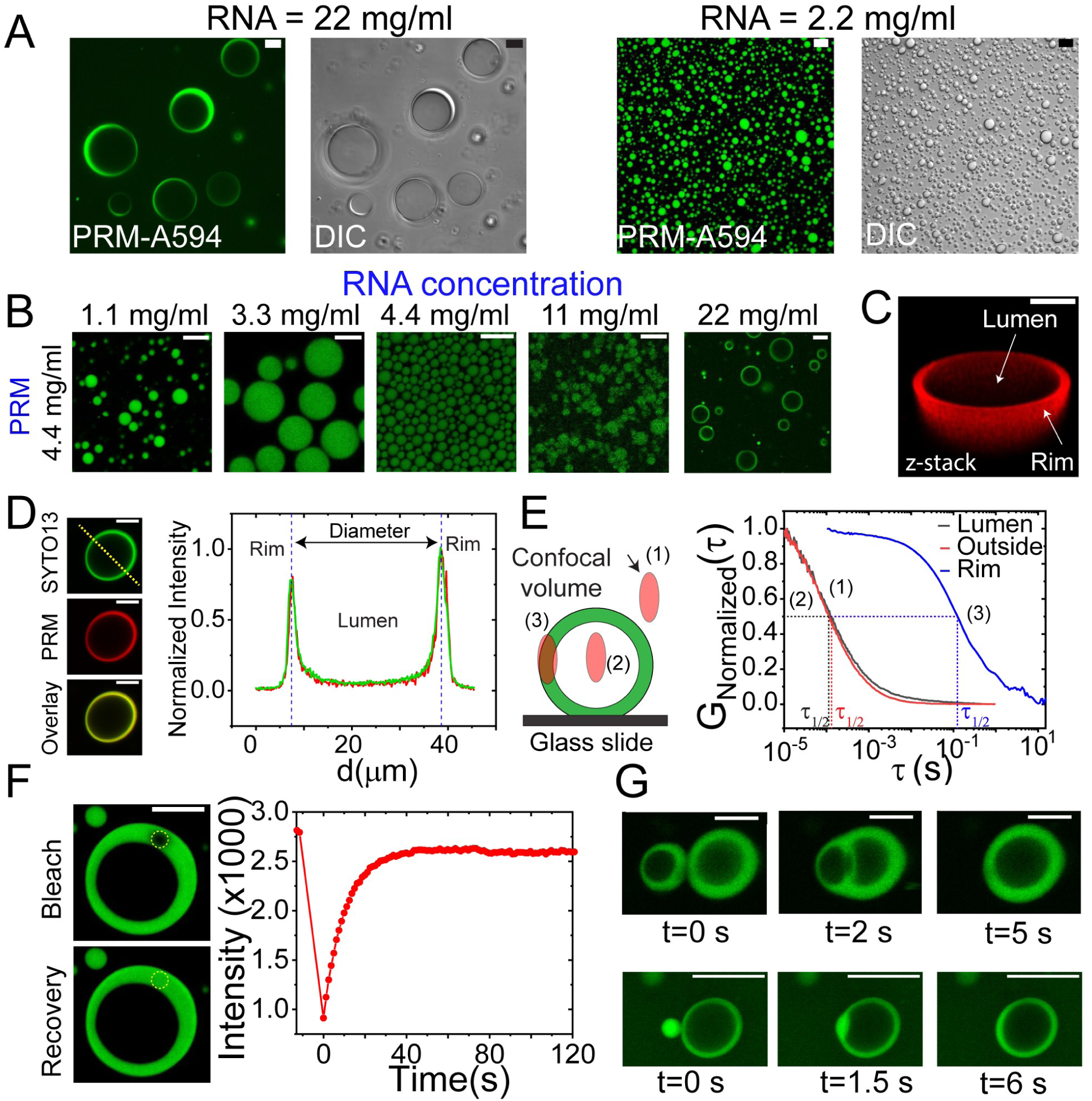
Protamine-RNA mixtures can form vesicular assemblies. ***(A)*** Fluorescence and DIC micrographs of PRM-RNA vesicles formed at 4.4 mg/ml PRM and 22 mg/ml poly(U) RNA (*i.e*., [PRM] = 5 *×*[RNA]; *left* panels). Contrastingly, at a lower PRM-to-RNA ratio, PRM-RNA mixtures form isotropic liquid droplets. Shown in *right* panels are micrographs of samples prepared upon mixing 4.4 mg/ml PRM and 2.2 mg/ml poly(U) RNA (*i.e*., [PRM] = 0.5 *×* [RNA]). ***(B)*** Fluorescence micrographs showing that mixture composition governs PRM-RNA droplets and vesicles formation. PRM concentration was fixed at 4.4 mg/ml. ***(C)*** 3D reconstruction of a PRM-RNA vesicle. ***(D)*** RNA (probed using SYTO13) and PRM (probed using Alexa594-labeled PRM) localize within the vesicle rim while the lumen has a hollow appearance. Fluorescence intensity of the lumen is similar to the external dilute phase. ***(E)*** Fluorescence correlation spectroscopy (FCS) autocorrelation curves for freely diffusing TMR-labeled Dextran-4.4k in three regions: inside the lumen, within the rim, and outside the hollow condensate. This experiment was done by focusing the confocal volume inside a hollow condensate, outside and on the rim (see *SI Appendix*, Fig. S6 & methods section). These autocorrelation curves suggest macromolecular diffusion is significantly slowed down (autocorrelation decays at ∼100 ms) within the rim as compared to the lumen and the external dilute phase (autocorrelation decays at ∼100 µs). ***(F)*** FRAP images and the corresponding intensity time-trace for PRM-A594 showing near complete fluorescence recovery of the hollow condensate rim. The yellow circle indicates the bleaching region (also see Supplementary Movie 1). ***(G)*** Optical tweezer-controlled fusion of two PRM-RNA hollow condensates (*top* panels) and a PRM-RNA droplet with a hollow condensate (*bottom* panels) (see also corresponding Supplementary Movies 2 and 3, respectively). Experiments were performed in 25 mM Tris-HCl buffer (pH 7.5). Fluorescent probe concentrations were ≤ 1% of the unlabeled protein and RNA. Fluorescence microscopy was performed with Alexa594-labeled PRM unless otherwise noted. Scale bars represent 10 µm.

Membranes are usually dynamic. For example, in lipid membranes, molecules diffuse within the membrane rim as a two-dimensional fluid (23). This observation led us to ask whether PRM-RNA vesicle rims also exhibit similar liquid-like properties. Our FCS measurements suggest that molecules are highly dynamic within the rim (Fig. 1*E*). To confirm this rapid internal dynamics within membranes, we utilized two orthogonal assays. *First*, we performed Fluorescence Recovery after Photobleaching (FRAP) experiment wherein we bleached a circular spot within the vesicle rim. FRAP recovery profile showed that the bleached region recovered to ∼100% fluorescence intensity within 30 seconds (Fig. 1*F*, Movie S1), suggesting a high molecular diffusivity within the hollow condensate’s rim. *Second*, we used optical tweezers to perform controlled fusion of two hollow condensates. Trap-controlled fusion experiments revealed that these hollow condensates fuse to initially form multi-compartment vesicular structures that rapidly relax into vesicles with single internal lumen (Fig. 1*G top* panels, Movie S2). This behavior is analogous to fusion of lipid membranes (24, 25). Passive fusion of vesicles was also observed during the initial stages of sample equilibration (*SI Appendix*, Fig. S7). Furthermore, we noticed that the hollow condensates can also fuse with small PRM-RNA spherical droplets, suggesting that they contain physicochemically compatible condensed phases (Fig. 1*G bottom* panels, Movie S3). This is further supported by comparing the FRAP recovery profiles for hollow condensates with the same for co-existing small PRM-RNA droplets (see *SI Appendix*, Fig. S8). Thus, similar to lipids within lipid-bound vesicle rims, protein and RNA molecules show a high degree of mobility within nucleoprotein-RNA membranes. Overall, our results clearly demonstrate that nucleoprotein-RNA complexes can form stable vesicular condensates that are structurally similar to lipid vesicles (see *SI Appendix*, Figs. S2, S4 & S5).

### Vesicular structures represent distinct condensed phases of protamine-RNA complexes

To determine the conditions at which PRM-RNA hollow condensates are stabilized, we mapped their thermodynamic state diagram as a function of mixture composition utilizing optical microscopy and solution turbidity measurements. Briefly, we recorded bright-field and fluorescence micrographs of PRM-RNA samples at a desired composition to determine whether the sample forms *droplets, hollow condensates*, or *a homogeneous mixture*. These measurements are summarized as a state diagram in Figure 2*A* (see also *SI Appendix*, Fig. S9*A*). This state diagram shows that PRM-RNA condensation is *reentrant* (see also *SI Appendix*, Fig. S9*B*) with a window-like two-phase coexistence region (19, 20). *In addition to the RNA-excess regime, we observed that hollow condensates appear at PRM-excess regime* (open blue circles in Fig. 2*A*). Inspection of the state diagram superimposed on the net charge concentration (estimated as *Q*_*polycation*_*C*_*polycation*_ − *Q*_*polyanion*_*C*_*polyanion*_; plotted as a color gradient in Figure 2*A*) reveals that stable hollow condensates are formed when there is a substantially excess net charge in the PRM-RNA mixture, regardless of whether this charge is negative (RNA excess) or positive (PRM excess). Electrophoretic mobility measurements indeed confirmed that *at RNA excess conditions, PRM-RNA complexes are negatively charged* and *at protein excess conditions, these complexes are positively charged* (Fig. 2*B* and *SI Appendix*, Fig. S10). Our fluorescence-based molecular partitioning assay suggests that, similar to vesicles under RNA-excess conditions, vesicles under PRM-excess regime also enrich both PRM and RNA within their rims whereas their internal lumen remains relatively depleted of both PRM and RNA (Fig. 2*C*).

**Figure 2:**
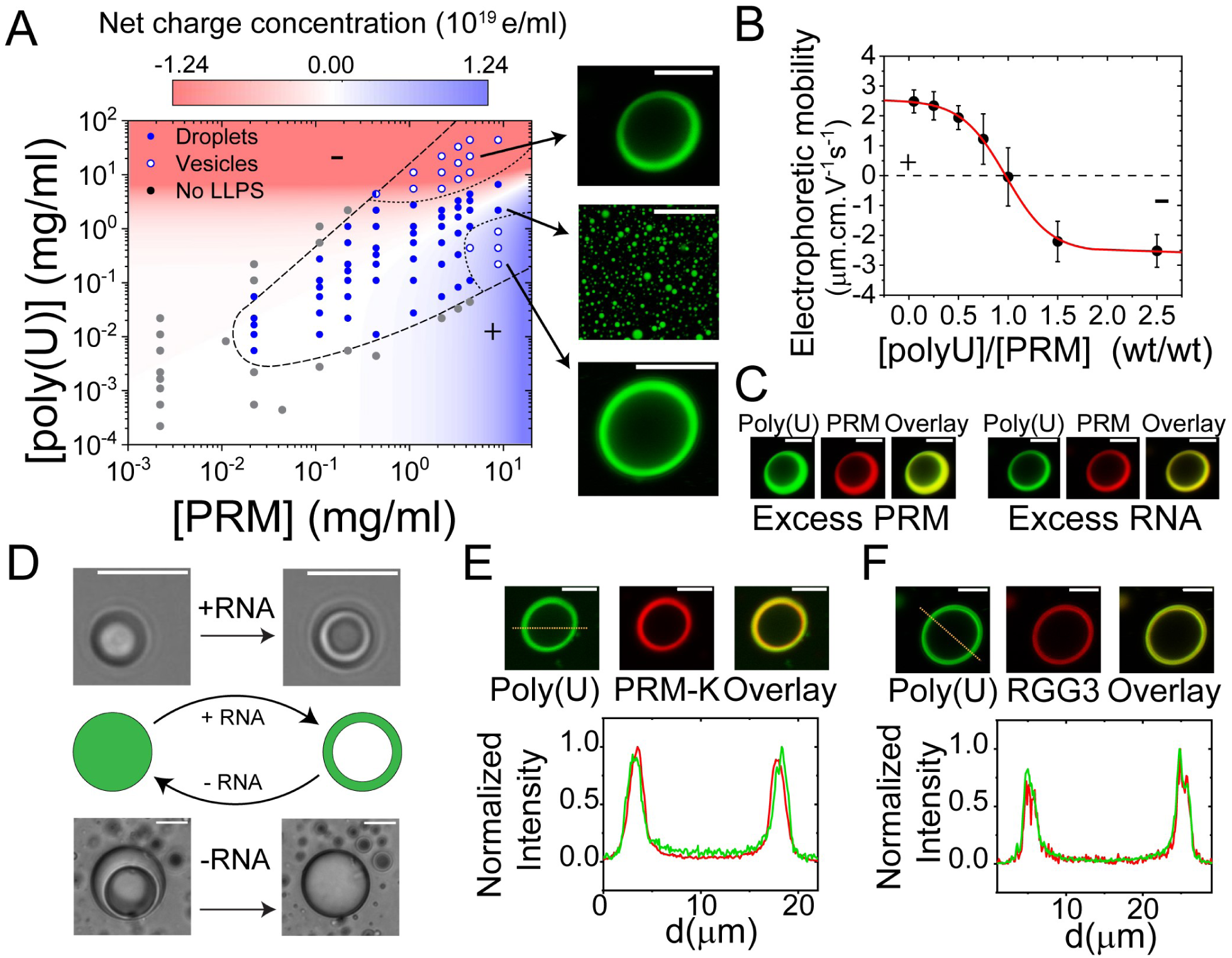
Vesicle-like polypeptide-RNA condensate is a thermodynamically stable phase. **(*A*)** Thermodynamic state diagram of PRM-poly(U) mixture shows three distinct phases: a homogeneous state (filled gray circles), PRM-poly(U) isotropic liquid droplets (filled blue circles), and PRM-poly(U) vesicles (open blue circles). The dashed line represents the boundary between homogeneous and phase separated regimes. The background colored shade represents the estimated concentration of electric charge (calculated from mixture composition). Hollow condensates were present at two narrow regimes within this state diagram, as indicated by dotted lines. The dashed and dotted lines are drawn as guides to the eye. Condensate imaging was done using PRM-A594. ***(B)*** Electrophoretic mobility measured by dynamic light scattering of PRM-poly(U) condensates as a function of poly(U)-to-PRM ratio. This data clearly shows charge inversion: condensates have a net positive charge at low RNA-to-PRM ratio and a net negative charge at high RNA-to-PRM ratio. PRM concentration was 1.1 mg/ml for this experiment. ***(C)*** Fluorescence micrographs of hollow condensates at RNA excess (upper edge of the LLPS region in ***A***) and PRM excess (lower edge of the LLPS region in ***A***) conditions. The excess PRM sample was prepared at 8.8 mg/ml PRM and 0.44 mg/ml poly(U). The excess RNA sample was made at 4.4 mg/ml PRM and 22 mg/ml poly(U). ***(D)*** A scheme showing droplet-to-vesicle and vesicle-to-droplet transformations upon RNA influx and removal, respectively (*middle* panels). DIC images of a PRM-RNA droplet [0.88 mg/ml PRM, 0.44 mg/ml poly(U)] transitioning to a vesicle upon RNA influx (*top* panels; also see *SI Appendix* Fig. S11 for experimental details; Supplementary Movie 4). DIC images of a PRM-RNA vesicle transitioning to a homogeneous droplet as a result of RNA removal (RNase-A treatment; *bottom* panels; see also Supplementary Movie 5). Fluorescence micrographs (*top* panels) and corresponding intensity profiles (*bottom* panels) of **(*E*)** hollow condensates formed by PRM-K (4.4 mg/ml) and poly(U) RNA (22 mg/ml), and ***(F)*** FUS^RGG3^ (4.0 mg/ml) and poly(U) RNA (20 mg/ml). Scale bars are 10 µm.

Next, we hypothesized that if these nucleoprotein vesicles indeed represent stable mesoscale structures in the state diagram, then influx or efflux of one of the components should be sufficient to reversibly induce droplet-to-vesicle transformations (Fig. 2*D*). To test this idea, we prepared and flowed PRM-RNA droplets [composed of 0.88 mg/ml PRM and 0.44 mg/ml poly(U)] into a microfluidic flow cell (*SI Appendix*, Fig. S11*A*). Then, we employed a dual-trap optical tweezer to trap two droplets and transported them to a second channel that was initially filled with buffer and connected to a poly(U) RNA inlet [poly(U) concentration = 10 mg/ml] (*SI Appendix*, Fig. S11*B*). This poly(U) concentration is higher than RNA concentration required for a droplet-to-vesicle structural transition under the experimental conditions. Once the flow of RNA started, we observed that the *PRM-RNA droplets rapidly transitioned to vesicles* (Fig. 2*D*; *SI Appendix*, Fig. S11*C*; Movie S4), whereas a control experiment with just the experimental buffer (lacking RNA) flow failed to produce a similar effect (*SI Appendix*, Fig. S11*C*). Contrastingly, when we removed RNA (partial) from preformed PRM-RNA vesicles by RNase-A treatment, *PRM-RNA vesicles rapidly transitioned to droplets* (Fig. 2*D*; Movie S5). These results clearly indicate that nucleoprotein condensates can dynamically undergo structural transition between vesicles and droplets via influx and removal of RNAs.

### Both arginine and lysine-rich polypeptides form hollow condensates with RNA

The RNA binding ability of protamine is primarily due to the 21 arginine residues (accounting for ∼ 63% of the protein sequence, *SI Appendix*, Fig. S1*A*). Previously, we and others reported that despite carrying the same amount of charge (+1e), arginine side chains are distinct compared to lysine as they are uniquely capable of mediating a hierarchy of ionic, cation-π, and π-π interactions with RNA (19, 26-30). To probe whether multi-modal Arg-RNA interactions are required for hollow condensate formation, we first tested whether a lysine variant of protamine (PRM-K; all 21 arginine residues are mutated to lysine, *SI Appendix*, Fig. S1*B*) can also form similar vesicular structures. Confocal microscopy imaging revealed a robust formation of stable hollow condensates for PRM-K under similar mixture compositions (Fig. 2*E*). Therefore, the ability to form vesicle-like assemblies appears to be independent of the identity of the positively charged residues (*i.e*., R *vs*. K) in nucleoproteins.

Both PRM and PRM-K have a high charge density [net charge per residue (NPCR) = 0.63]. To test whether high charge density is a prerequisite for vesicular condensate formation, we next used another naturally occurring disordered RNA-binding polypeptide, the RGG3 domain of FUS (FUS^472-504^), which is similar in length to PRM but only contain 7 arginine residues and no lysine residues (NPCR = 0.2, see *SI Appendix*, Fig. S1*C*). Similar to the PRM system, FUS^RGG3^ also formed hollow condensates at excess RNA conditions (Fig. 2*F*). In all these systems, both proteins and RNAs showed co-localization in the rim, similar to PRM-RNA vesicle membranes. Together, these results highlight the generality of vesicular structure formation in mixtures of disordered RNA-binding polypeptides and RNAs. Based on the previous studies on condensate physical properties for arginine-to-lysine variants of R/G-rich proteins (19, 26), we speculate that polypeptide primary sequence will tune the compositions at which vesicular assemblies are stable as well as the physicochemical properties of the vesicle rim. However, as discussed in the following section, the ability of these systems to spontaneously form hollow condensates appears to be generic at disproportionate mixture compositions and is linked to the heterotypic nature of this associative phase separation.

### Anisotropic protein-RNA complexes drive hollow condensate formation

Our state diagram analysis of PRM-RNA mixture clearly indicate that vesicular structures are formed at distinct mixture composition and concentration regimes (Fig. 2*A*). Many theories of weak and strong polyelectrolyte complexation have predicted the existence of micellar and lamellar phases (31-35), but none of these theories predict hollow condensate formation as a function of the mixture composition. To conceptualize this phenomenon, we turn to the reentrant phase transition model by Shklovskii and Zhang (36). This model postulates that at conditions far from the equal stoichiometry, mixtures of a long polyanion and a short polycation will form a partially condensed complex with a *tadpole-like* geometry consisting of a neutral head and a charged tail (Fig. 3*A*; *top* panels). The tadpole head can be considered as a nano-condensate. We propose that, above a threshold concentration (*SI Appendix*, Note-1), heads of neighboring tadpoles may coalesce to form small spherical condensates (referred to as *micelles* henceforth), where the condensate surfaces are decorated with the bare segments of RNA chains (Fig. 3*A*; *middle* panel; *SI Appendix*, Figs. S12*A*). This condensation of protein-bound segments of the RNA chains can be driven by differential intra-complex solvation since the free part of an RNA chain remain charged, and therefore, are expected to have higher effective solvation volume than the condensed segment of the chain (37). Once formed, although these micellar condensates are fluid-like (*SI Appendix*, Fig. S12B), they are stabilized against coalescence-driven growth due to their low interfacial energy (*SI Appendix*, Fig. S12). We hypothesize that this is due their surfaces being decorated by the part of RNA chains that remain free (37). Upon increase in the number density of micellar condensates, the total free energy of the mixture increases due to excluded volume interactions between micelles, triggering a micelle-to-vesicle transition (see *SI Appendix*, Note-1). The vesicular condensates provide an additional interface allowing redistribution of bare RNA chains within the vesicle lumen, thereby reducing the free energy of the system (Fig. 3*A bottom* panels).

**Figure 3:**
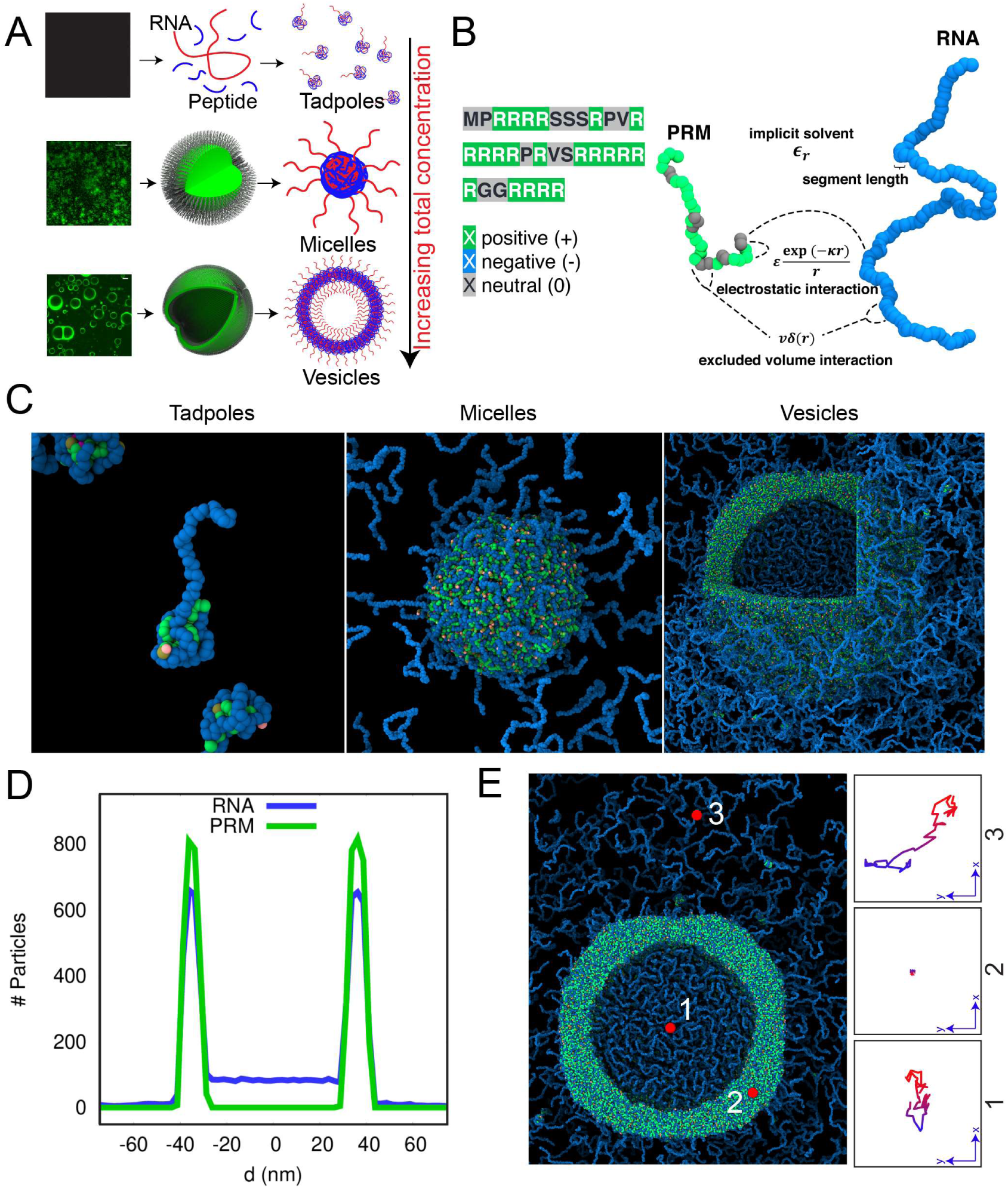
Intra-complex disproportionation drives the formation of vesicular assemblies. ***(A)*** A scheme for the formation of RNA-protein hollow condensates. *Left column panels* show representative experimental observations (fluorescence micrographs). The proposed mechanism is shown in right two columns. At low concentrations, nucleoprotein-RNA mixtures form tadpole-like complexes. Increasing the total concentration leads to the formation of small spherical micellar assemblies. At relatively high total concentration, nucleoprotein-RNA vesicular structures are formed. ***(B)*** Schematic representation of the protein and RNA chains and the interaction potential employed in our molecular dynamics (MD) simulations. ***(C)*** Equilibrium MD configurations showing tadpole, micelle, and vesicle formed under charge disproportionate conditions, CRNA = 5 *×*CPRM. (Also see *SI Appendix*, Figure S13). (***D***) Density profiles of PRM and RNA chains for the vesicle obtained from the MD trajectory. (See *SI Appendix*, Figure S5 for experimental data) ***(E)*** Diffusion trajectories of a tagged particle (red sphere) located within the lumen, within the rim, and outside of the vesicle. (Also see *SI Appendix*, Figure S13).

Although the vesicle formation in RNA-protein mixtures can be qualitatively explained based on the Shklovskii-Zhang model for tadpoles, the model does not predict vesicle formation. To test whether tadpole-like complexes can self-assemble into micellar and vesicular structures, we performed molecular dynamics (MD) simulations using coarse-grained models of PRM and RNA chains. Amino acid residues and nucleic acid bases in these chains were represented as single beads (Fig. 3*B*). The interatomic potential employed in our simulations contains bonded, electrostatic, and short-range pairwise interaction terms as shown in Figure 3*B*. The simulation setup and the coarse-grained model employed is discussed in *SI Appendix, Materials and Methods*. The MD simulations were performed with *C*_*RNA*_ *= 5 × C*_*PRM*_ at three different RNA concentrations (0.01 mg/ml, 7.4 mg/ml, and 15.80 mg/ml) that mimic the three experimental conditions of low, intermediate, and high concentration regimes. The system size ranges from 10^3^-10^6^ particles across the studied concentrations. The representative equilibrium structures sampled from MD simulations at these three concentrations are shown in Figure 3*C*. In our MD simulations, we find that compositionally disproportionate mixtures indeed form tadpole-like structures at low chain volume fraction, micellar condensates (with the free RNA decorating the condensate surface) at intermediate volume fraction, and hollow vesicle-like condensates at relatively high volume fraction (Fig. 3*C, SI Appendix*, Fig. S13 & Movie S6). Therefore, we infer that a vesicle-like phase is intrinsically accessible to protein-RNA mixtures at high concentrations and disproportionate mixture compositions. The simulation-derived density profiles of the PRM and RNA chains in the hollow vesicle-like structure is shown in Figure 3*D* and is comparable to the experimental density profile shown in Figure 1*D*. To compare the relative diffusivities in various phases, we tag a single particle on the outside, the lumen, and the rim of the vesicle and follow its motion over a 10 ns (four orders of magnitude in coarse-grained simulation time-scale) segment of MD trajectory (Fig. 3*E, SI Appendix*, Fig. S14). We observe that the particles in the lumen and outside the vesicle have comparable diffusivities as they diffuse over a significant volume, indicating a low-density phase. Contrastingly, the particle within the rim has a much slower diffusivity and is confined close to its initial position, indicating a condensed phase. These results are consistent with our experimental FCS autocorrelation curves shown in Figure 1*E* and *SI appendix*, Fig. S6. Furthermore, similar to our microfluidics experiments (Fig. 2*D*), our simulations reveal that increasing the RNA concentration in the external dilute phase leads to a transition from a spherical micelle to a hollow condensate (*SI Appendix*, Fig. S15 & Movie S7).

Analogous to the excess RNA simulations, we also performed MD simulations with *C*_*PRM*_ *= 5 × C*_*RNA*_ at three different PRM concentrations (0.01 mg/ml, 7.0 mg/ml, and 14.60 mg/ml). We find that akin to excess-RNA conditions, the system equilibrates to tadpole-like structures at low concentration, micellar condensates at intermediate concentration, and hollow vesicle-like structures at high concentration in excess PRM regimes (*SI Appendix*, Fig. S16*A*). We also observe that the vesicle has high density of PRM and RNA chains at the rims and has a low PRM concentration in its lumen (*SI Appendix*, Fig. S16*B*). This is similar to the density profile observed in vesicles formed at excess RNA conditions (Fig. 3*D*).

### Both biological and synthetic heterotypic systems can form hollow condensates at disproportionate mixture compositions

Our experimental observations and MD simulation results collectively suggest that vesicle formation might be an intrinsic feature of ternary mixtures that undergo liquid-liquid phase separation driven by heterotypic electrostatic interactions. To experimentally verify this idea, we tested the ability to form vesicles by several pairs of oppositely charged biological and synthetic polyelectrolytes. First, we kept the polycation (PRM) unchanged and changed the identity of the polyanion. We observed that PRM can form vesicle-like hollow condensates with inorganic polyphosphate (polyP), polyglutamic acid [poly(E)], and poly(acrylic acid) (PAA) at similar composition and concentration regimes (Fig. 4*A*-*C, SI Appendix*, Fig. S17). Second, we changed the polycation, keeping the polyanionic RNA unchanged. Similar to the PRM-poly(U) system, we observe that the RNA can also form hollow condensates upon interaction with a synthetic cationic polyelectrolyte, poly(Allylamine) (PAH) under similar conditions (Fig. 4*D, SI Appendix*, Fig. S17). We also changed identities of both polycation and polyanion, and observed that PAH-polyP mixture form vesicles at disproportionate mixture compositions (Fig. 4*E, SI Appendix*, Fig. S17). Finally, we tested whether cellular RNA, a mixture that contains both structured and unstructured RNAs, can form vesicular assemblies. Indeed, we observed that total RNA from *Saccharomyces cerevisiae* can form stable micron-sized hollow condensates upon interaction with an R/G-rich polypeptide at excess RNA condition (Fig. 4*F, SI Appendix*, Fig. S17). These results reveal that vesicular assemblies represent a distinct yet generic condensed phase in biological as well as synthetic heterotypic coacervate-forming systems.

**Figure 4.**
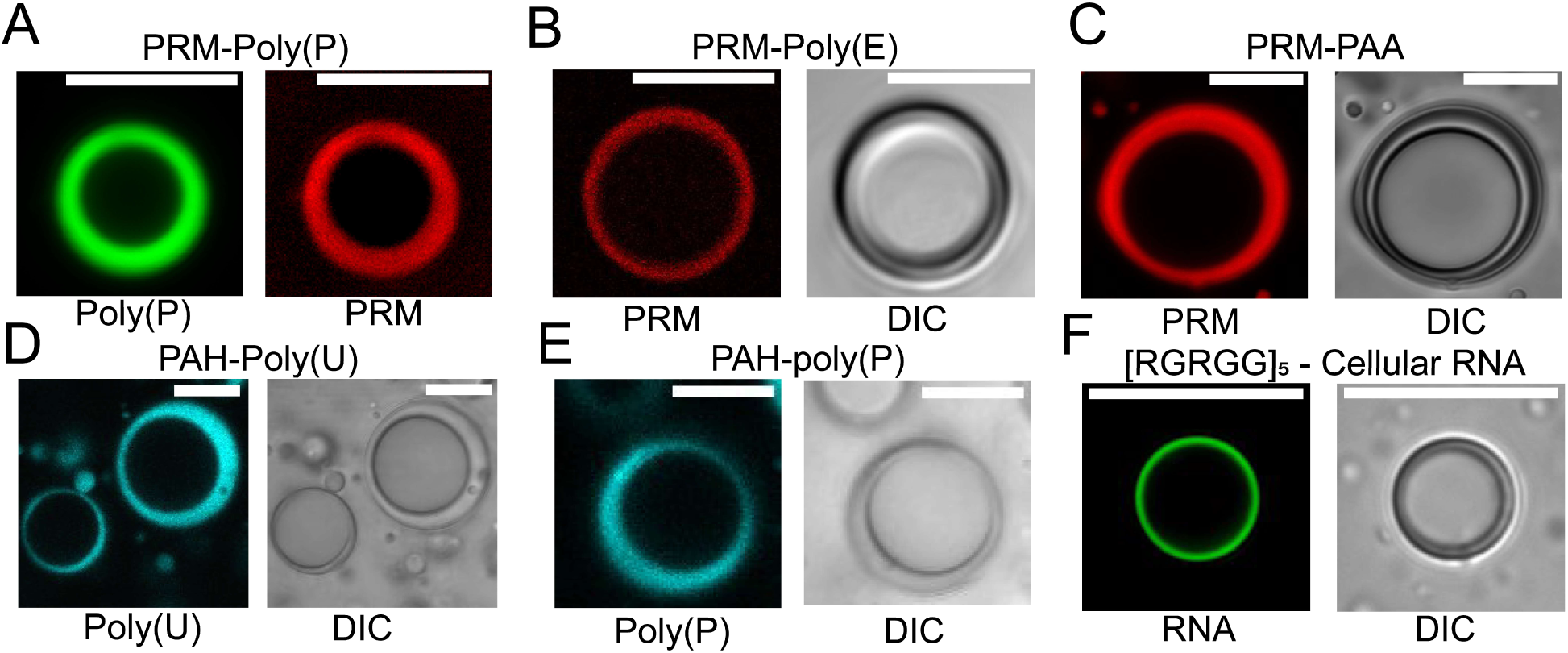
Hollow condensates are widely observed in various biological and synthetic ternary systems. Fluorescence and DIC micrographs of hollow condensates formed by PRM (4.4 mg/ml) and polyP (22.0 mg/ml, ***A***); PRM (17.6 mg/ml) and poly(E) (18 mg/ml, ***B***); PRM (8.8 mg/ml) and poly(acrylic acid) (PAA; 22 mg/ml, ***C***); PAH (40 mg/ml) and poly(U) (4.0 mg/ml, ***D***); PAH (70 mg/ml) and polyP (4 mg/ml, ***E***); and [RGRGG]5 (0.024 mg/ml) and cellular RNA (8.9 mg/ml, ***F***). See also *SI Appendix* Figure S17. Scale bars represent 10 µm.

### Protein-RNA hollow condensates exhibit molecular ordering, size-dependent permeability, and selective encapsulation

Oppositely charged disordered biopolymers primarily attract each other through electric monopoles that are typically isotropic, *i.e*., that they lack spatial directionality (38). However, in the case of amphiphiles, hydrophobic tail groups are packed together in the bilayer interior, which may give rise to spatial ordering. In the mesoscale, many amphiphilic assemblies, such as vesicles, are therefore characterized by a liquid crystalline ordering (39, 40). Our MD simulation results, however, suggest that the rims of the protein-RNA vesicles contain alternating bands of positively charged and negatively charged residues interspersed with the neutral residues of the PRM chains (*SI Appendix*, Fig. S16*C* and *D*). Computed structure factors of hollow condensates from our MD simulation indicate a radial arrangement of RNA chains that is absent in the case of spherical micelles (SI Appendix, Fig. S18). This organization of protein and RNA chains may result in local molecular ordering in protein-RNA vesicle membranes, akin to classical lipid-bound membranes. To test the existence of molecular ordering experimentally, we imaged the PRM-RNA vesicles utilizing polarization light microscopy. Interestingly, we observed that PRM-RNA hollow condensates exhibited signatures of liquid-crystalline ordering on the rim (41) (Fig. 5*A*), whereas PRM-poly(U) droplets did not show any signature of ordering under similar conditions (*SI Appendix*, Fig. S19*A*). To check whether the ordering is specific to PRM-RNA vesicles, we recorded polarization micrographs of PRM-K–RNA, FUS^RGG3^-RNA, and PRM-polyP vesicles (*SI Appendix*, Fig. S19*B*-*D*). All these vesicles demonstrated an optical birefringent pattern, which is characteristic of molecular ordering in the rim (41). Therefore, our cross-polarization light microscopy images of hollow condensates suggest that molecular ordering in the rim is generic to protein-RNA vesicles, as is the case for many amphipathic lipid-bound membranes.

**Figure 5:**
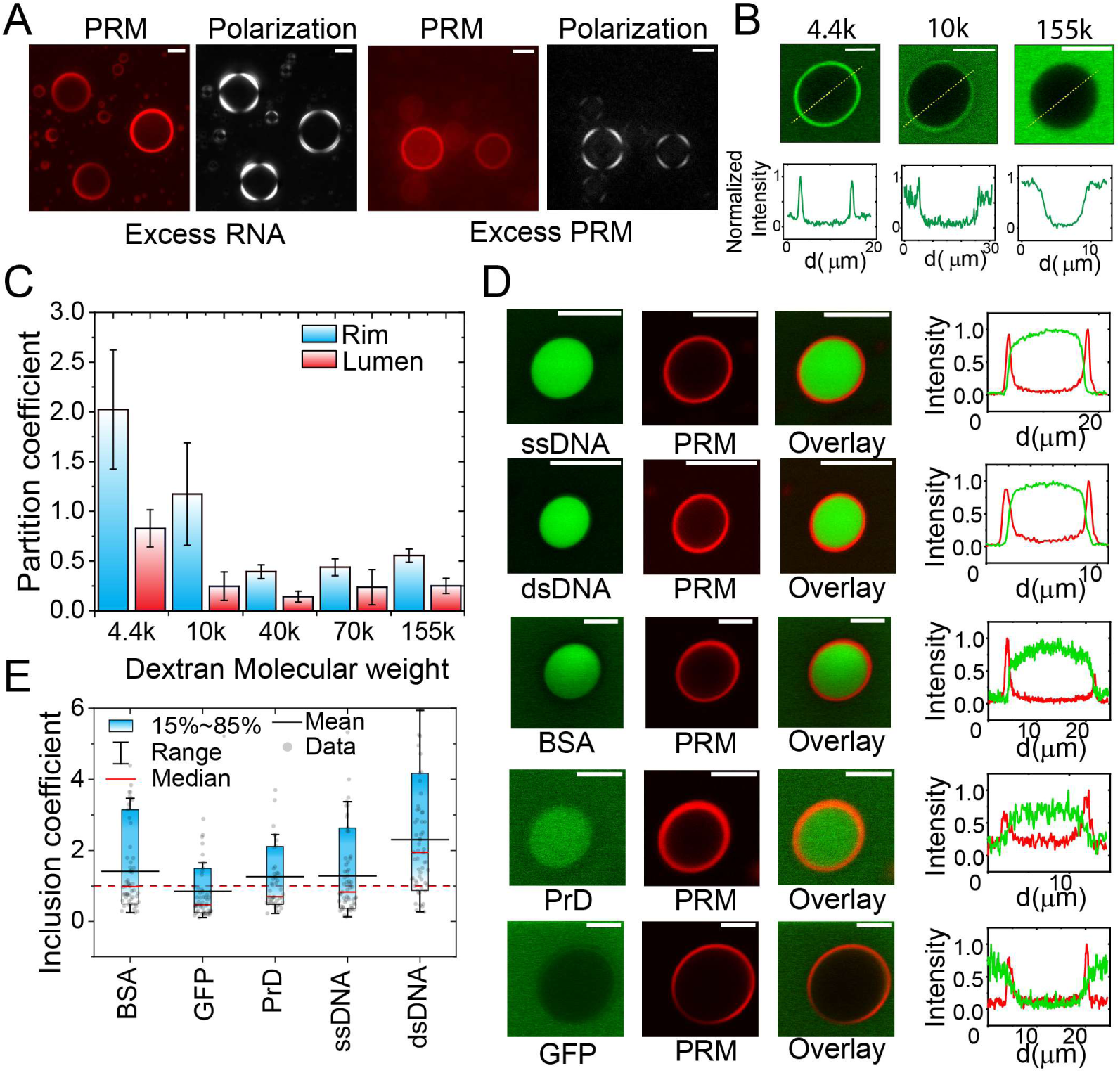
PRM-RNA hollow condensate rims exhibit classical lipid-bound membrane-like properties. ***(A)*** Optical images of PRM-RNA hollow condensates with cross-polarizing light show birefringence, indicating molecular ordering in the vesicle rims. Corresponding fluorescence micrographs are also shown. ***(B)*** Fluorescence images and corresponding intensity profiles of TMR-labeled dextran probes of different molecular weights reveal size-dependent partitioning in PRM-RNA hollow condensates. The low molecular weight dextran, Dex-4.4k, partitions to the rim whereas the high molecular weight dextran, Dex-155k, remains excluded from the rim. See also *SI Appendix* Figure S20. ***(C)*** A bar plot of size-dependent partitioning data of dextrans into the rim and lumen of PRM-RNA hollow condensates. ***(D)*** Partitioning of nucleic acids and proteins into PRM-RNA vesicles. The intensity profiles are measured along a horizontal line passing through the center of individual hollow condensates. *BSA*: Bovine Serum Albumin; *GFP*: Green Fluorescent Protein; *PrD*: Prion-like domain of FUS. ***(E)*** Inclusion coefficients (defined as the ratio of *mean intensity in the lumen* to *mean intensity in the external dilute phase*) are shown. The statistics were estimated using a minimum of 50 different hollow condensates per sample. Individual points are shown as gray filled circles. Scale bars are 10 µm.

Our data indicate that physicochemical properties of the nucleoprotein vesicle rim are similar to nucleoprotein droplets (Fig. 1*F-G*). Recently, it was reported that ribonucleoprotein droplets can act as a size-dependent filter allowing permeability of molecules that are smaller than the condensate mesh-size (42). To test whether protein-RNA vesicle membranes have similar size-dependent permeability, we added fluorescently labeled dextran probes of various sizes to pre-formed hollow condensates. We observed that Dextran-4.4k (MW 4400 Da) partitioned favorably within the rim, whereas a larger dextran, Dextran-10k, weakly partitioned to the rim. Further increase in the dextran size resulted in probe exclusion from both the rim and the lumen (Fig. 5*B, SI Appendix* Fig. S20). A summary of these results is presented in Figure 5*C*, where we plotted the partition coefficients in the rim/lumen for dextran probes of different molecular weights ranging from 4.4k to 155k. These experiments suggest that, similar to ribonucleoprotein droplets, PRM-RNA hollow condensate rims act as a size-dependent filter. Based on the hydrodynamic radius of Dextran-10k in an aqueous buffer (42), we estimate the mesh-size of PRM-RNA vesicles to be ∼ 2.3 nm.

A major application of amphiphilic vesicles is in cargo trafficking and delivery, which require entrapment of the cargo molecules within the internal vesicular lumen. We, therefore, tested the ability of protein-RNA vesicles to encapsulate a wide variety of client biomolecules (*e.g*., proteins and nucleic acids). Using two-color confocal imaging, we determined partition coefficient for each client (defined as *I*_*lumen*_*/I*_*outside*_). We observed that both ssDNA and dsDNA are enriched within the lumens of PRM-RNA vesicles (Fig. 5*D*). For proteins, we observed that biomolecule loading into PRM-RNA vesicle is protein-specific. For example, bovine serum albumin (BSA) and the prion domain of FUS (PrD) were enriched in the PRM-RNA vesicle lumen whereas the green fluorescent protein (GFP) remains excluded. These results suggest that the PRM-RNA vesicles selectively enrich biomolecules and indicate a potential utility of these vesicles as stimuli responsive dynamic cargo carriers.

## Conclusion

Heterotypic interactions between R/K-rich LCDs and RNAs have been shown to be crucial for the biomolecular condensation of many nucleoproteins (8, 26, 43-47). Previously, we showed that RNA has a stoichiometry-dependent effect on the LLPS of R/K-rich LCDs [*i.e*., at low RNA levels, LLPS of R/K-rich LCDs is facilitated and at high RNA levels, their LLPS is inhibited (18, 19)]. Here, we show that partially condensed nucleoprotein-RNA complexes can form distinct stable supramolecular topologies, such as tadpoles, micelles and vesicles. Our observed nucleoprotein vesicles bear significant structural and functional similarities with lipid vesicles. For example, similar to lipid membranes, nucleoprotein-RNA vesicle membranes exhibit molecular ordering and is able to selectively encapsulate biomolecules. However, unlike lipid vesicles, nucleoprotein vesicles remain highly dynamic and can readily undergo reversible *vesicle-to-droplet* phase transition in a stimuli responsive fashion. Thus, nucleoprotein vesicles may represent a dynamic mode of creating multilayered membrane-less assemblies that are ubiquitous in the subcellular space. Interestingly, the estimated *in vivo* RNA concentration in the mammalian cell nucleus [∼ 8.5 mg/ml, (48)] is comparable to the *in vitro* RNA concentrations that led to stable vesicle formation in this study [11 mg/ml poly(U) and 8.9 mg/ml cellular RNA with PRM and FUS^RGG5^, respectively, Figs. 2*A* & 4*F*)].

Physicochemically, the rims of nucleoprotein vesicles are similar to nucleoprotein droplets. As in the droplets, biomolecules in the rim are mobile and the rims of two vesicles in contact can undergo fusion. We also observe that the formation of vesicles is not specific to nucleoprotein-RNA complexes, rather is more generic to ternary systems that undergo LLPS via predominantly heterotypic electrostatic interactions. This observation is puzzling in light of the known principles of vesicle-like assembly formation, which suggest that the geometric anisotropy of individual building blocks is a key driver for the “bottom-up” self-assembly of vesicular structures (49). For example, lipids and lipid-inspired di-block copolymers that are known to form vesicles feature anisotropic architecture with two distinct domains, which vastly differ either in their solvent interactions or in their intermolecular inter-domain interactions (49, 50). Furthermore, protein vesicles that have been reported in the literature are formed by engineered biopolymers with similar di-block architecture (51-53). Low-complexity biological and synthetic polymers utilized in our current study, however, are intrinsically isotropic in isolation (*i.e*., they do not have blocks of distinct physical and chemical properties). Based on our MD simulation, we propose that partially complexed isotropic polypeptide and/or RNA chains can form anisotropic building blocks (such as tadpoles) that may lead to vesicular topologies. We speculate that the formation of hollow condensates may be a general phenomenon for multi-component systems of associative polymers which show a liquid phase transition via obligate heterotypic interactions. This opens up a new avenue to design and fabricate coacervate-based supramolecular assemblies with tunable multi-layered topologies.

We envision several key implications of our observations. *First*, nucleoprotein condensates can attain multilayered topologies via the vesicle formation pathway. This pathway is orthogonal to the recently proposed mechanism of multiphasic condensate formation via coexisting liquid droplets in a multi-component mixture (5). *Second*, stimuli-responsive sequestration through the dynamic formation/dissolution of vesicles of low-complexity polymers may have been utilized by prebiotic and protobiotic systems. Third, by coupling with stronger physical and/or chemical cross-linking strategies, protein-RNA vesicles can be utilized as stimuli responsive lipid-free cargo delivery systems for biotechnological applications (*e.g*., drug/gene delivery, insecticide/pesticide release). These cargo delivery systems can be formulated carrier-free by directly utilizing proteins or nucleic acids of interest and therefore, achieve a high degree of target loading. *Finally*, these dynamic assemblies can be utilized to fabricate novel stimuli-responsive microscale systems.

## Acknowledgements

The authors gratefully acknowledge UB north campus confocal imaging facility (supported by National Science Foundation MRI Grant: DBI 0923133) and its director, Mr. Alan Siegel for helpful assistance. The authors acknowledge Dr. Evan Nelsen (Lumicks) for his help with PSF characterization. The authors also acknowledge helpful discussions with Ms. Taranpreet Kaur, Dr. George Thurston, and Dr. Wei Wang at various stages of manuscript preparation.

## Funding

We gratefully acknowledge support for this work from University at Buffalo, SUNY, College of Arts and Sciences to P.R.B. and funding from the National Institute on Aging (NIA) of the National Institutes of Health (R21 AG064258) to P.R.B.

## Author contributions

P.R.B. and M.M.M. conceived the idea. I.A. and P.R.B. designed the experiments. I.A. preformed the experiments and analyzed the data with help from P.R.B. M.R. and D.P. designed and performed the MD simulation. P.R.B., M.M.M., and I.A. wrote the manuscript with input from M.R. and D.P.

## Conflict of interest

P.R.B, M.M.M., and I.A. have a pending patent application related to the present study: “Lipid-free polyionic vesicles and methods of making and using same, provisional application US 62/958,039 filed by University at Buffalo, SUNY, January 2020.

