## Supplementary Appendix for "Phase Transition of RNA-protein Complexes into Ordered Hollow Condensates"

**This PDF file includes:**

Supplementary text
Figures S1 to S26
Legends for Movies S1 to S7
SI References

**Other supplementary materials for this manuscript include the following:**

Movies S1 to S7

### Materials and Methods

**Protein/RNA samples and labeling.** Salmon protamine (P4005) was purchased from Sigma-Aldrich and used without any further purification. The third RGG box of FUS (RGG3; AA: 472-505), protamine-lysine variant (PRM-K with 21 R-to-K substitutions) and $[RGRGG]_5$ were synthesized by GenScript USA Inc. (NJ, USA; ≥ 90% purity) and used without further purification. All synthesized polypeptides have a C-terminal cysteine for site specific labeling. Polyuridylic acid [poly(U); molecular weight = 600-1000 kDa, LOT#: 028M4144V & 089M4052V] and yeast total RNA was purchased from Sigma-Aldrich. Protein and RNA stock preparation and protein labeling was done as described in earlier work (1). Poly(U) RNA was re-suspended in RNase Free Water at concentrations ≥ 80 mg/ml, then aliquots were made and stored at −20 °C for later use. Given poly(U) RNA is known to undergo phase separation independent of proteins(2), the undiluted stocks were checked using a bright-field microscope to ensure aggregate-free RNA in our experiments. SYTO13, Pierce recombinant Green Fluorescent Protein (GFP) and albumin from bovine serum, fluorescein conjugate (BSA) were purchased from Thermo Fisher Scientific Inc. Prion-like domain of FUS (PrD; AA: 1-173) was expressed, purified and labeled as described in our earlier work (3). Polyphosphate (p100, medium chain; MW ~ 11.5 kDa) was purchased from Kerafast, Inc. Poly(L-Glutamic Acid) was purchased from Alamanda Polymers (MW ~120 kDa). Pluronic F-127, poly(allylamine) solution (20 wt. % in $H_2O$, MW ~ 15 kDa), poly(Acrylic acid) (MW ~ 1.8 kDa) and Tetramethylrhodamine (TMR) conjugated Dextrans of various molecular weights (4.4 kDa, 10 kDa, 40 kDa, 70 kDa, and 155 kDa) and DAPI were purchased from Sigma-Aldrich. Fluorescently labeled ssDNA and dsDNA (ssDNA sequence: TGCAGATTGCGCAATCTGCA; dsDNA sequence: 5'-TGCAGATTGCGCAATCTGCA-3') was purchased from IDT. Custom synthesized FAM-RNA ([6FAM]UGAAGGAC) was purchased from Sigma-Aldrich.

**Fluorescence correlation spectroscopy (FCS).** Samples were prepared at 4.4 mg/ml protamine concentration and 22 mg/ml poly(U) RNA. Dextran-TMR (MW: 4400 Da) or Alexa594-PRM was added to the buffer prior to mixing PRM and poly(U) (*i.e.* before forming the hollow condensates). Samples were then injected into a Pluronic coated (1% wt/vol) 25 mm x 75 mm x 0.1 mm single chamber custom-made flow-cell. Photon arrival times were collected for 3 minutes at room temperature (23 ± 1 °C) using a laser scanning confocal microscope (LUMICKS C-Trap) with a single-photon Avalanche photodiode (sAPD) at 100 MHz acquisition rate. This experiment was done by conducting a point scan and focusing the microscope on the desired region of interest. For the Lumen, the microscope was focused on the center of the vesicle and the photon count was observed to be similar to that of the external dilute phase. However, we saw a multifold increase in the photon count when focusing on the rim (see Fig. S6*B*), indicating that the signal is coming from the hollow condensate's rim. We also characterized the PSF of the microscope to ensure the accuracy of the obtained data (see Fig. S6*C*). Photon arrival times were then autocorrelated using pycorrelate python library (see pycorrelate documentation for the algorithm (4), *version 0.2.1*). For each case in Figure 1E, 5 autocorrelation curves were averaged and plotted using OriginPro 2018b software.

**Fluorescence Recovery After Photobleaching (FRAP).** FRAP experiments were done using a Zeiss LSM 710 laser scanning confocal microscope, equipped with a 63× oil immersion objective (Plan-Apochromat 63×/1.4 oil DIC M27). Samples were prepared using 4.4 mg/ml PRM and 22 mg/ml poly(U) with ~1% Alexa594 labeled protamine, and then placed on a Pluronic-coated Nunc Lab-Tek chambered coverglass (ThermoFisher Scientific Inc.) at room temperature (23 ± 1 °C). A circular region of interest (ROI) was bleached with 100% laser power for 2-5 seconds and fluorescence intensity from the ROI was subsequently collected until recovery was complete. For analysis, FRAP recovery traces were fitted with a single exponential function unless otherwise noted.

**Controlled fusion using optical traps.** Two condensates were trapped using identical 1064 nm trapping lasers (LUMICKS, C-Trap, 60× water-immersion objective). Due to the low refractive index mismatch, the laser power was increased to prevent loss of condensates from the traps. One trap was moved at a constant velocity of ~1 μm/s towards the other trap to initiate condensate fusion. Coalescence movies were collected and processed thereafter using Fiji-ImageJ software (v1.52p) (5). For two hollow condensate fusion, the sample was prepared at 4.4 mg/ml PRM and 22 mg/ml polyU in a 15 mM Tris-HCl (pH 7.5) 100 mM NaCl buffer. For droplet fusion with a hollow condensate, the sample was prepared at the same concentrations in a 15 mM Tris-HCl (pH 7.5) buffer.

**Confocal imaging, partition measurements and mesh size determination.** Samples were prepared at 4.4 mg/ml protamine concentration and 22 mg/ml poly(U) RNA in a 15 mM Tris-HCl buffer (pH 7.5). Fluorescent probes were added directly to the buffer at 0.1%-1% labeled-to-unlabeled protein ratio. Samples were inserted into a Pluronic-coated (Pluronic 1% wt/vol) 25 mm x 75 mm x 0.1 mm single chamber custom-made flow-cell. Multicolor fluorescence images were collected using a laser scanning confocal microscope (LUMICKS, C-Trap, 60× water-immersion objective). Partition coefficients were calculated as the ratio between the intensity in the lumen and the intensity in the external dilute phase. For the partition coefficients of ssDNA, dsDNA, GFP, BSA and PrD, ~50 Hollow condensates were analyzed to ensure statistical accuracy. For estimating the condensate mesh size, dextran probes were added *after* forming the hollow condensates and equilibrated for 15 minutes prior to imaging. For time and temperature series (Fig. S2), a Zeiss laser scanning confocal microscope (LSM 710, 63× oil immersion objective (Plan-Apochromat 63×/1.4 oil DIC M27)) was used with a temperature controlled stage (Pecon temperature incubator with a stage-top heating insert).

**Polarization light microscopy.** Samples were prepared in a buffer containing 15 mM Tris-HCl (pH 7.5) and placed on a Pluronic-coated (Pluronic 1% wt/vol) 25 mm x 75 mm x 0.1 mm (concentrations are indicated in corresponding figure legends). Condensates were imaged using a Leica DMi 8 inverted microscope equipped with sCMOS Leica camera (20x and 100x Objectives were used). Polarized LED white light was produced using Polaroid dichroic filters.

**Thermodynamic state diagram.** The protein-RNA state diagram was determined using optical microscopy. Protein-RNA samples were prepared in a buffer containing 5 mM Tris-HCl (pH 7.5). Each sample was inspected for 2 minutes under a bright field microscope (Zeiss

Primo-vert inverted iLED microscope with 40× objective lens and a Zeiss Axiocam 503 monochrome camera) for condensate formation. This was repeated for several RNA-to-protein stoichiometry at different initial protein concentrations. OriginPro 2018b software was used to plot the resulting phase diagram.

**Electrophoretic light scattering.** Electrophoretic mobility of protein-RNA complexes was measured using a dynamic light scattering (DLS) setup (ZetasizerNano ZS; Malvern Instruments Ltd.) utilizing M3-PALS (Phase Analysis Light Scattering) method. Samples were prepared at 100, 250 and 500 µM protamine concentrations and various poly(U) concentrations as indicated in the respective plots (buffer: 25 mM Tris-HCl, pH 7.5).

**Solution Turbidity measurements:** Polypeptide and RNA mixtures were prepared at desired peptide concentration with variable RNA concentrations. The buffer contained 25 mM Tris-HCl (pH 7.5). The absorbance was measured at 350 nm using a NanoDrop oneC UV-Vis spectrophotometer at room temperature after ~100 seconds of sample equilibration with a 1.0 mm optical path length. Turbidity plot was generated via gradual RNA titration. Measurements were performed in triplicates. The phase boundary curve was obtained by plotting the data using OriginPro software.

**RNA jump experiments utilizing microfluidics.** Droplets formed by 0.88 mg/ml protamine and 0.44 mg/ml poly(U) RNA were flowed into a single channel within a custom-made microfluidics chip (see Fig. S11*A*) using an automated pressure-box (LUMICKS, u-Flux). A second channel was filled with 10 mg/ml poly(U) RNA, which is appropriate for protamine-RNA vesicle formation for 0.88 mg/ml protamine. The flow rate was kept at minimum to ensure laminar flow and avoid mixing of the channels. Two droplets were trapped in the first channel using 1064 nm lasers (LUMICKS, C-Trap) with minimal power and transported into the second channel (see Fig. S11*B*). The droplet-to-vesicle transition was imaged using a 1.31 Mpix microscope camera (Sensor Type: CMOS mono, Framerate: 60 fps, LUMICKS, C-Trap). A corresponding control experiment was run with the second channel filled with experimental buffer. The control experiment revealed no detectable changes in droplet morphology within the timeframe of our experiments (30 minutes).

**RNase-mediated vesicle-to-droplet transition.** Vesicle samples were prepared at protamine concentration of 4.4 mg/ml and poly(U) concentration of 22 mg/ml in 15 mM Tris-HCl (pH 7.5). The vesicles were imaged with a bright field microscope (Zeiss Primo-vert inverted iLED microscope with 40× objective lens and a Zeiss Axiocam 503 monochrome camera), followed by addition of RNase-A (Thermo Scientific) to the sample at a final concentration of 0.057 mg/ml. The mixture then was continuously imaged. A corresponding control experiment was also performed where the same volume of experimental buffer was added (without RNase-A), which produced no detectable changes in vesicle morphology within the timeframe of our experiments (15 minutes).

**Molecular Dynamics simulation.** In this study, we have employed a single residue/base resolution coarse-grained polyelectrolyte model for protein and RNA chains. For the amino acids, we utilize the same coarse-grained parameters as those employed by Dignon *et al.* (6) to study phase behavior of intrinsically disordered proteins. The potential energy function contains bonded, electrostatic, and short-range pairwise interaction terms. Bonded interactions are modelled by a harmonic potential $k_r(r - r_0)^2$ with a spring constant $k_r = 10\ kJ/\text{Å}^2$ and an equilibrium bond length of $r_0 = 3.8\ \text{Å}$. Electrostatic interactions are modeled using a Coulombic term with Debye-Hückel electrostatic screening to account for salt concentration, having the functional form:

$$E_{ij}(r) = \frac{q_i q_j}{4\pi D r} exp(-\frac{r}{\kappa}) \qquad (1)$$

where $\kappa$ is the Debye screening length and D = 80, is the dielectic constant of the solvent (water). We set the Debye screening length $\kappa = 0.1$, which corresponds to the physiological buffer condition at room temperature. The RNA chain is modeled as a one bead per nucleotide model compatible with the protein model with the only difference being the addition harmonic angular term $k_\theta(\theta - \theta_0)^2$ to model the stiffness of the RNA chains, where spring constant $k_\theta = 1.0\ kJ$ and equilibrium angle $\theta_0 = 1.78$ rads. For the short-range pairwise interaction we employ a Lennard-Jones (LJ) 6-12 potential. The simple model consisting of Debye-Hückel electrostatics and LJ 6-12 potential considered in our study is able to capture and describe the experimental observations.

To mimic the experimental conditions at excess RNA conditions, the systems were initialized by placing PRM and RNA chains randomly in a cubic box with $Conc_{RNA} = 5 \times Conc_{PRM}$. To explore the transition from tadpole to micelle to vesicle structures, the simulations were performed at three different RNA concentrations; 0.01 mg/ml, 7.4 mg/ml and 15.80 mg/ml, respectively. We performed molecular dynamics (MD) simulations at 298 K using the Langevin thermostat with a friction coefficient of $\gamma = 0.01$ and time step of 0.01$\tau$. MD runs were performed on graphical processing units (GPUs) using the HOOMD-blue package v2.5.1(7, 8). The equilibrium run was performed for 2 $\times 10^6$ steps with an integration time step of 0.01$\tau$ and the production run was performed for an additional $2\times10^6\ \tau$ where $\tau$ is characteristic time-scale for the coarse-grained system. The number of protein and RNA chains and the corresponding box dimension is shown below.

| | # PRM chains | # RNA chains | Box dimension |
|---|---|---|---|
| Tadpole | 25 | 50 | 450 |
| Micelle | 200 | 400 | 100 |
| Vesicle | 5870 | 11740 | 240 |

**Table S1.** Number of protein and RNA chains employed in simulations.

**Diagrams and schemes.** All plots and graphs were produced using OriginPro software. Confocal images processing was done using Fiji-ImageJ. Diagrams and schemes and the final assembly of figures was done using Adobe Illustrator 2019 and Autodesk Autocad 2018. Molecular dynamics snapshots and movies were produced using VMD software v1.9.

**Supplementary Figures**

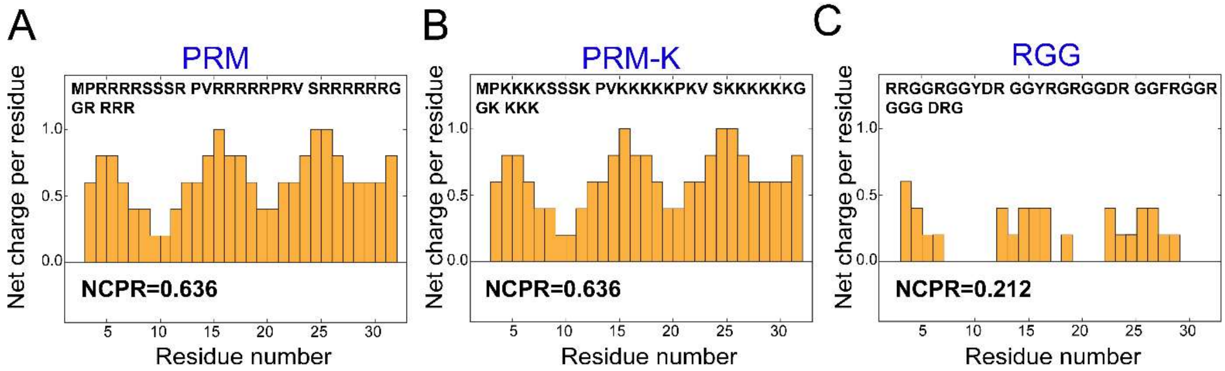


**Fig. S1.** Net charge per residue (NCPR), as calculated by the CIDER software (9), plotted against residue number for ***(A)*** Protamine (PRM), ***(B)*** Arg-to-Lys protamine variant (PRM-K), and ***(C)*** FUS RGG3 ($FUS^{472\text{-}504}$).

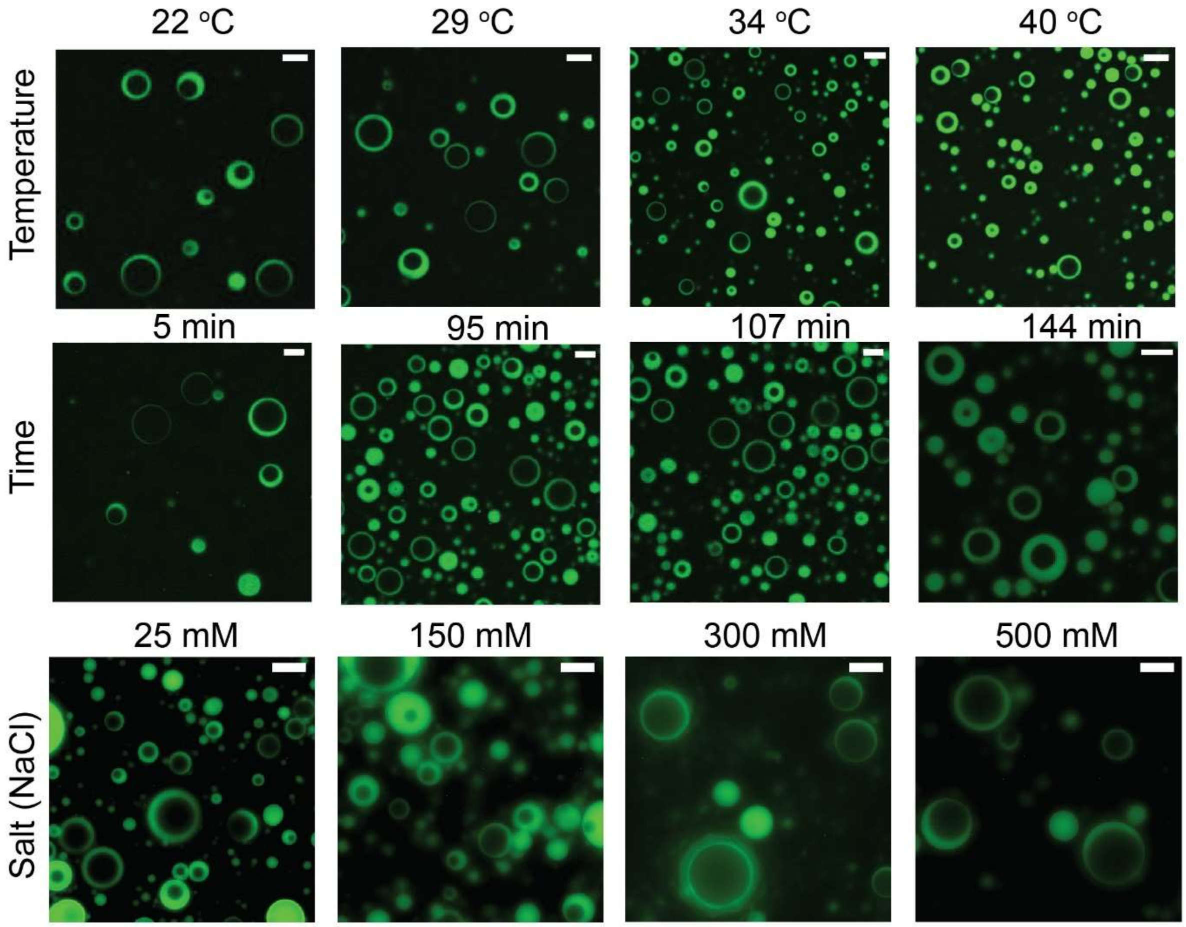


**Fig. S2.** Hollow condensates imaged at different temperatures (top row), time points (middle row), and salt concentrations (bottom row). Representative micrographs shown in each row are obtained from experimentally independent data sets. Sample contains 4.4 mg/ml PRM and 22 mg/ml poly(U) RNA in 25 mM Tris-HCl buffer, pH 7.5, unless otherwise indicated. Scale bars represent 10 μm. The temperature and time series were recorded using a confocal microscope (Zeiss LSM 710) whereas the variable salt images were obtained using an epifluorescence microscope (Zeiss primo-vert inverted iLED microscope, equipped with a Zeiss Axiocam 503 monochorme camera).

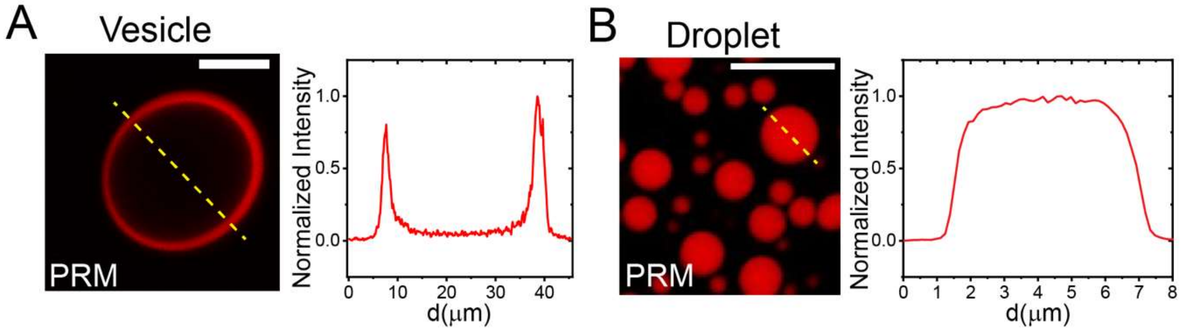


**Fig. S3.** Fluorescence micrographs and corresponding intensity profiles of a PRM-RNA vesicle [4.4 mg/ml PRM and 22 mg/ml poly(U), ***A***] and PRM-RNA droplet [4.4 mg/ml PRM and 2.2 mg/ml poly(U), ***B***]. Samples were prepared in 15 mM Tris-HCl buffer, pH 7.5. Scale bars represent 10 μm.

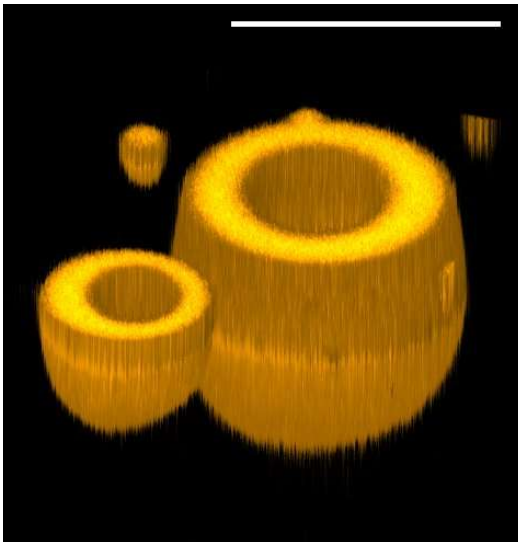

**Fig. S4.** 3D reconstitution of PRM-poly(U) vesicles from a Z-stack imaging experiment. Vesicle forming mixture composition: 4.4 mg/ml PRM and 22 mg/ml poly(U) in 15 mM Tris-HCl buffer, pH 7.5. Scale bar represents 10 μm.

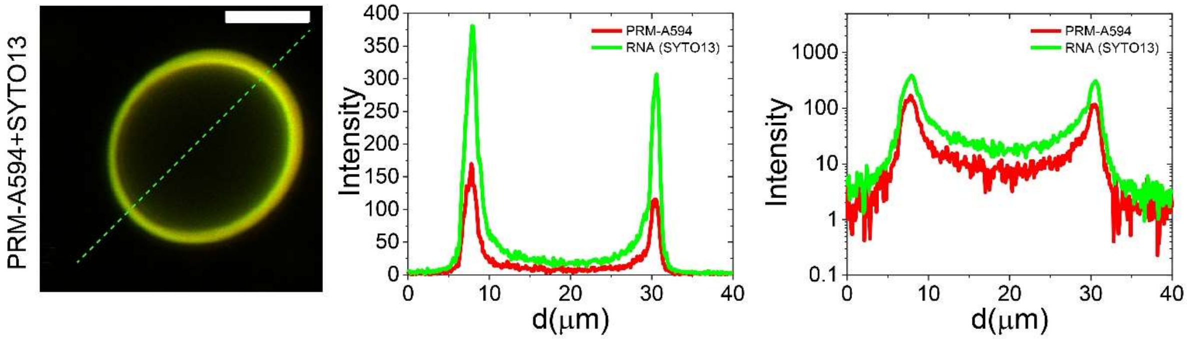


**Fig. S5.** Raw data for partitioning of PRM and RNA shown in Fig. 1*D*, *Main-text*. ***(left)*** Fluorescent image of PRM-poly(U) hollow condensate (4.4 mg/ml PRM and 22 mg/ml poly(U) in 15 mM Tris-HCl buffer (pH 7.5)). ***(middle)*** Intensity profiles for PRM-A594 and SYTO13 labeled poly(U). ***(right)*** Same profile plotted on a logarithmic scale (Y-axis) to better visualize the intensity variations across the hollow condensate. Scale bar represents 10 μm.

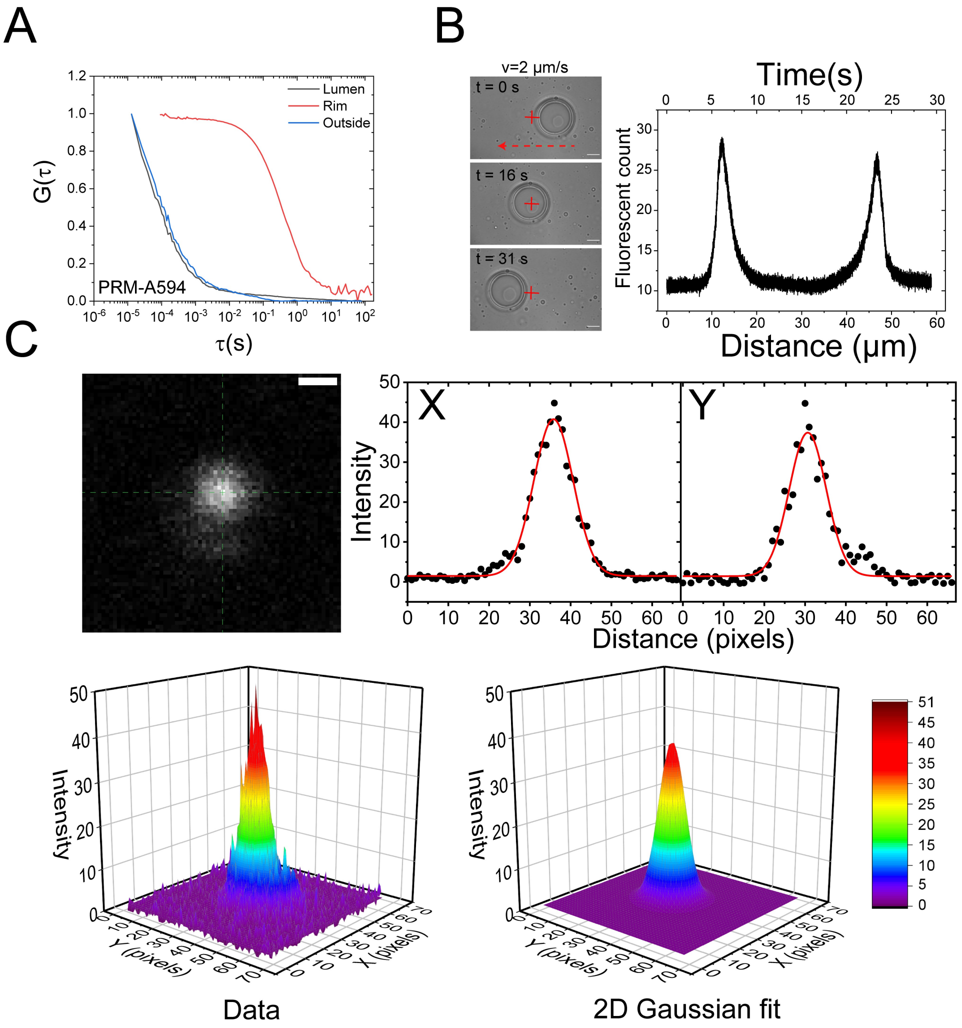


**Fig. S6.** ***(A)*** Fluorescence autocorrelation curves for the lumen, rim and the external dilute phase of a PRM-RNA hollow condensate. Here, Alexa594-PRM was utilized as the probe. ***(B)*** Photon count versus time for a hollow condensate as the stage is moved across the confocal volume (indicated by the red cross). The overall curve looks identical to the intensity profiles shown in Figure 1*F*, *Main-text*. Scale bar represents 10 μm. ***(C)*** Point spread function (PSF) for the confocal microscope used for FCS. The PSF was measured by scanning a 200 nm bead with a pixel size of 30 nm. Profiles across the maximum intensity in the X and Y directions are shown and were fitted using a Gaussian function. The full width at half maximum (FWHM) was found to be 341 nm for the X-direction and 320 nm for the Y direction. 2D Intensity plot of the raw data and the Gaussian fit are also shown. Scale bar represents 300 nm.

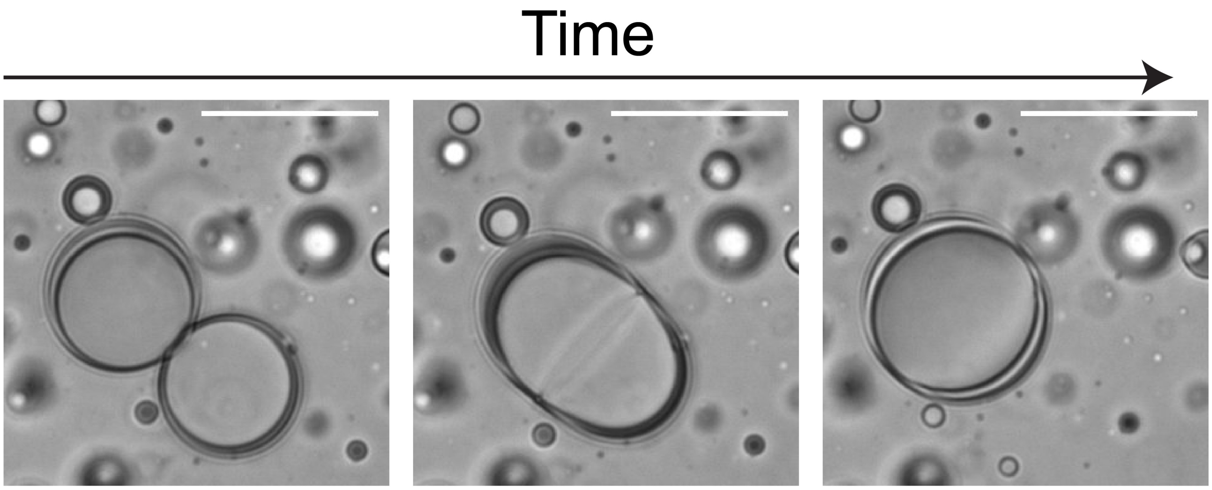


**Fig. S7.** Passive fusion of two hollow condensates (time-stamp: 0, 1.0 and 3.0 seconds from left to right panels). Sample contains 4.4 mg/ml PRM-K and 22 mg/ml poly(U) RNA in 15 mM Tris-HCl buffer, pH 7.5. Scale bars represent 10 μm.

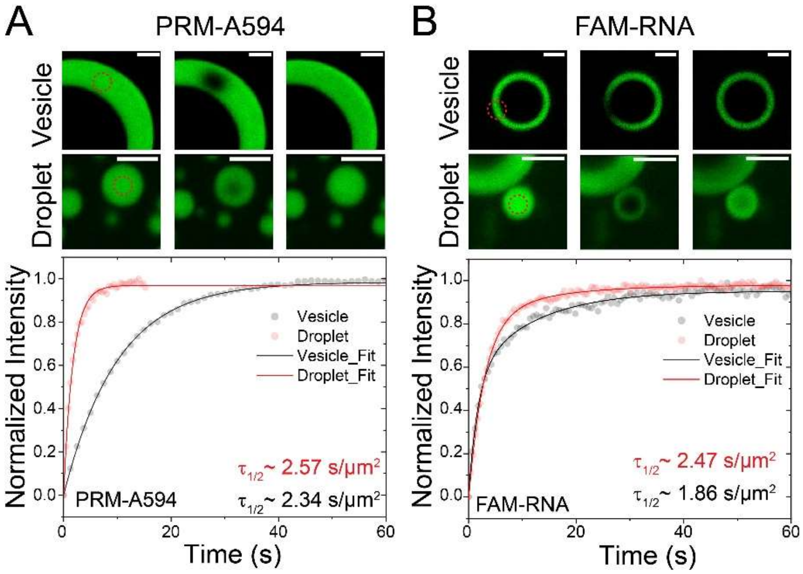


**Fig. S8.** FRAP recovery traces for ***(A)*** PRM-Alexa594 and ***(B)*** FAM-RNA (Fluorescein labeled UGAAGGAC) in a hollow condensate's rim and a co-existing small PRM-RNA droplet. Solid lines represent exponential fits to the data. For PRM-Alexa594, a single exponential fit was sufficient to give good fits to the data, while for FAM-RNA, we used a bi-exponential fit. We normalized the recovery half-time with the bleaching area to compare between the two scenarios. The estimated normalized half-times seem to be close to one another for the case of Alexa594-PRM but different in the case of FAM-RNA. We attribute this to a non-uniform bleaching in the case of vesicles since laser bleached a full segment of the vesicle and the diffusion was only in the direction adjacent to the vesicle rim. Scale bars represent 3 μm.

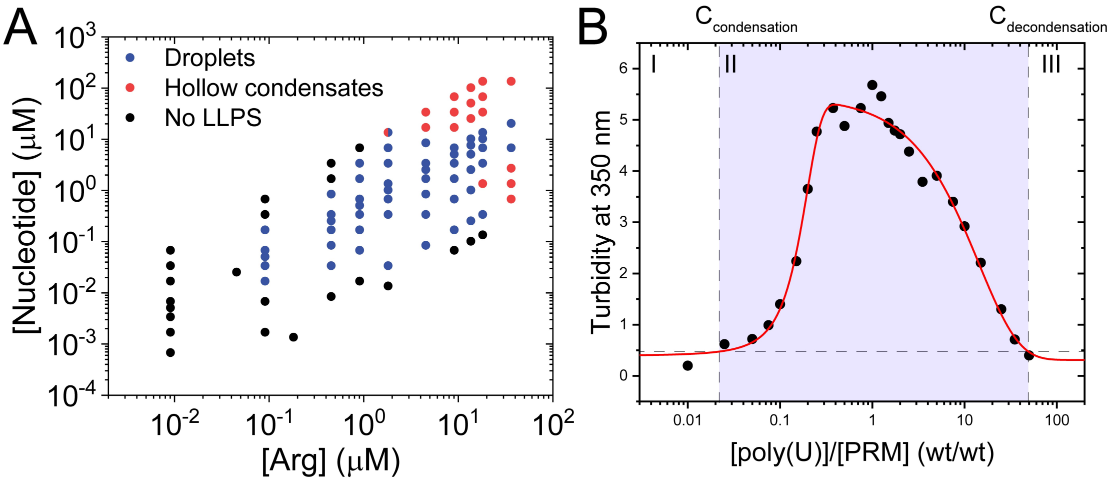


**Fig. S9.** ***(A)*** Thermodynamic state diagram for PRM-poly(U) mixtures. This is identical data to Figure 2*A*, *Main-text*, but plotted against arginine and nucleotide concentrations. ***(B)*** Solution turbidity at 350 nm as a function of poly(U)-to-PRM ratio. The horizontal line at 0.5 indicate the turbidity threshold for phase separation as judged by optical microscopy. The shaded region indicates the window of LLPS. PRM-poly(U) system undergoes condensation at ratios ≥ 0.02 and reentrant dissolution at ratios ≥ 42. The black filled circles are the data and the red line is a guide to the eye. PRM concentration was 0.33 mg/ml in 25 mM Tris-HCl buffer, pH 7.5.

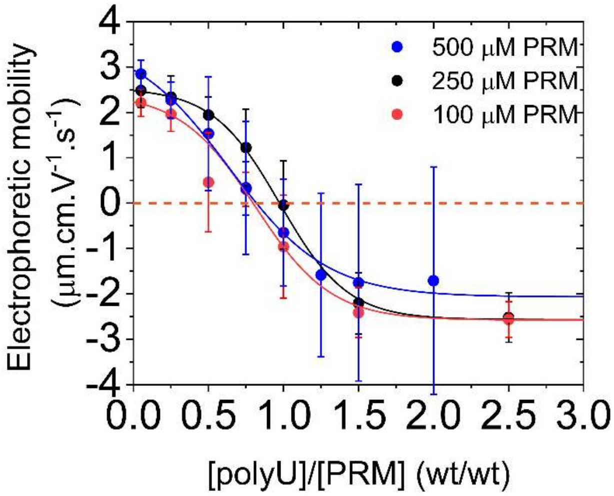


**Fig. S10.** Electrophoretic mobility measurement of PRM-poly(U) mixtures as a function of poly(U)-to-PRM ratios for three different PRM concentrations [100 μM PRM ≡ 0.44 mg/ml PRM]. At low RNA concentrations, complexes bear a positive charge while at high RNA-to-PRM ratio, complexes are negatively charged. Error bars represent the width of the mobility distribution of respective samples. For all concentrations, charge inversion is observed around a ratio of 1:1 (wt/wt).

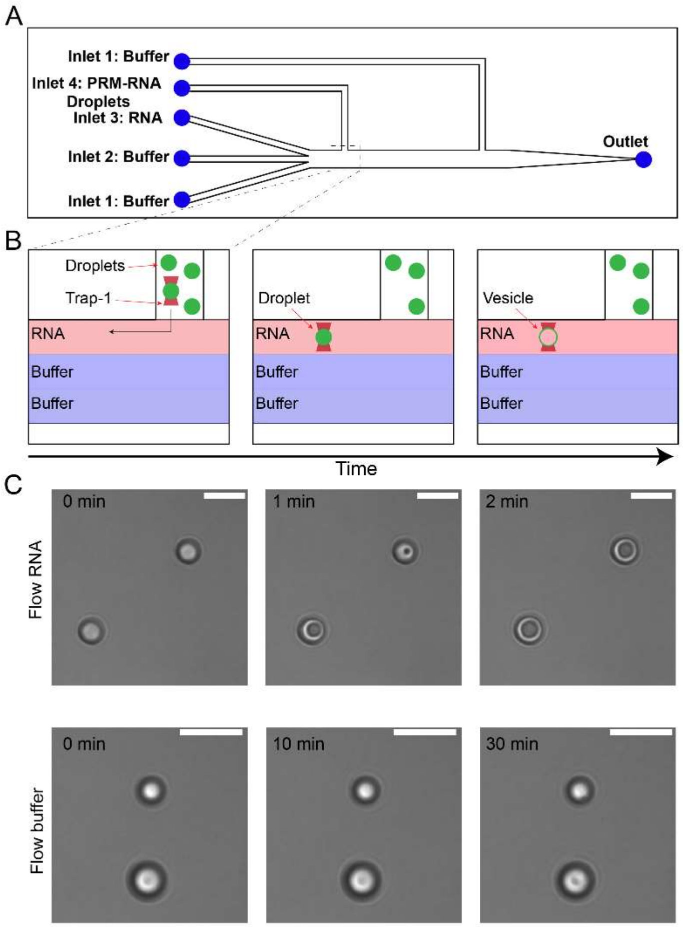


**Fig. S11.** A schematic depiction of ***(A)*** the microfluidics flow cell design, and ***(B)*** the experimental method employed. Here, droplets were flown in inlet 4 until they were observed in the channel. RNA and buffer were flown in inlets 1-3, as indicated. The flow was kept at minimal pressure to ensure constant laminar flow. The constant laminar flow ensures spatial separation of the fluids in different channels. A droplet was trapped in the droplet channel (channel 4) and moved to the RNA channel (channel 3). As RNAs interact the droplet, a droplet-to-vesicle transition was observed (***C***, *top* panels, see also Movie S4). When the same experiment was performed with buffer (without RNA) flown in channel 3, no visible change was observed for a period of 30 minutes (***C,*** *bottom* panels). Scale bar represents 10 µm.

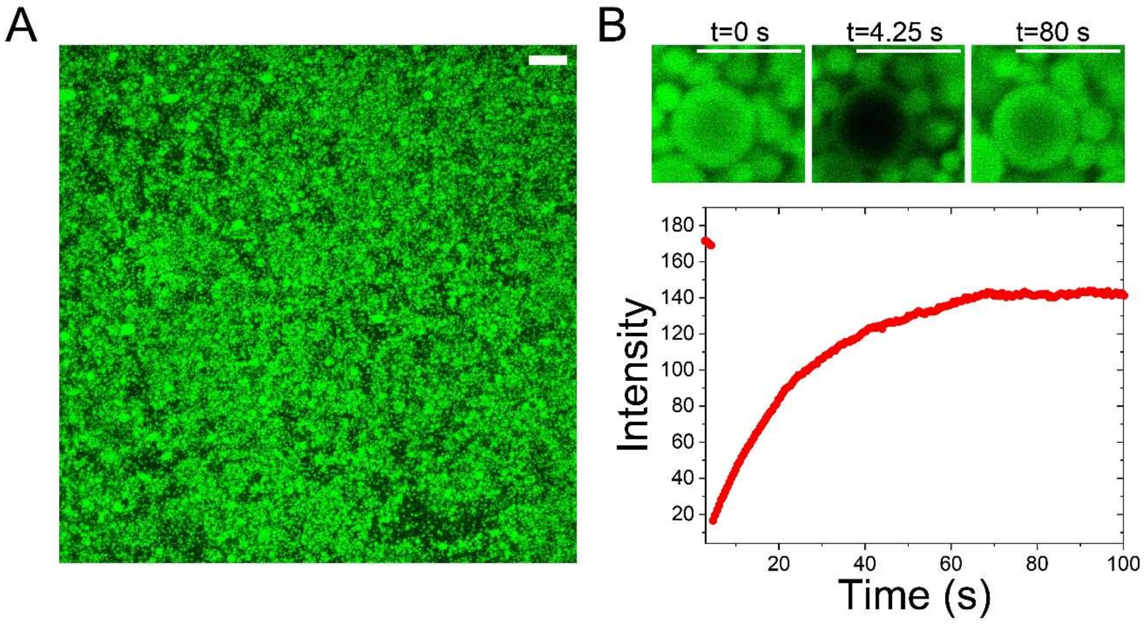


**Fig. S12.** ***(A)*** A representative fluorescence micrograph of PRM-poly(U) micelles formed at 2.2 mg/ml PRM and 12.8 mg/ml poly(U) in 25 mM Tris-HCl buffer (pH 7.5). Sample was imaged after 40 minutes. Scale bar represents 10 µm. ***(B)*** Fluorescence recovery after photobleaching (FRAP) of a micellar condensate formed at 2.2 mg/ml PRM and 12.8 mg/ml poly(U) in 25 mM Tris-HCl buffer. Scale bars represent 5 µm.

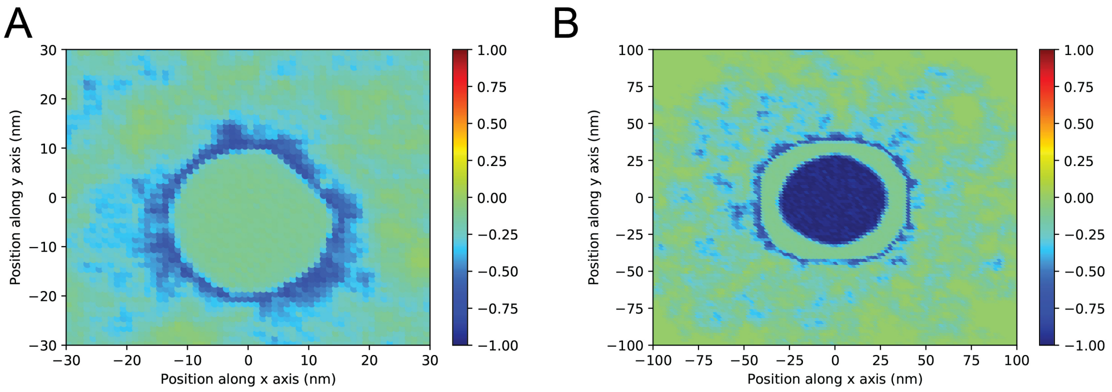


**Fig. S13.** Charge distribution of (***A***) micelle and (***B***) vesicle formed under excess RNA conditions. The charge distribution was calculated by dividing the simulation box into 1×1×1 nm$^3$ bins and computing the net charge in the bins using the final configuration at the end of the production run.

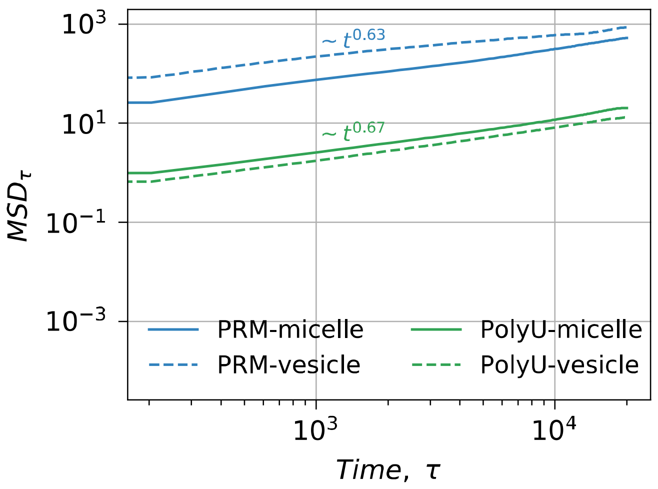


**Fig. S14.** Confined dynamics of RNA and protein monomers leads to sub-diffusive behavior**.** Mean square displacements of PRM and Poly(U) monomers as a function of simulation time $\tau$ computed during equilibrium sampling of micelles and vesicles respectively (for a period of 10 ns which is about four orders of magnitude larger than the simulation time step). Also see Fig. 3*C* in the *Main-text* for the structure of these vesicles.

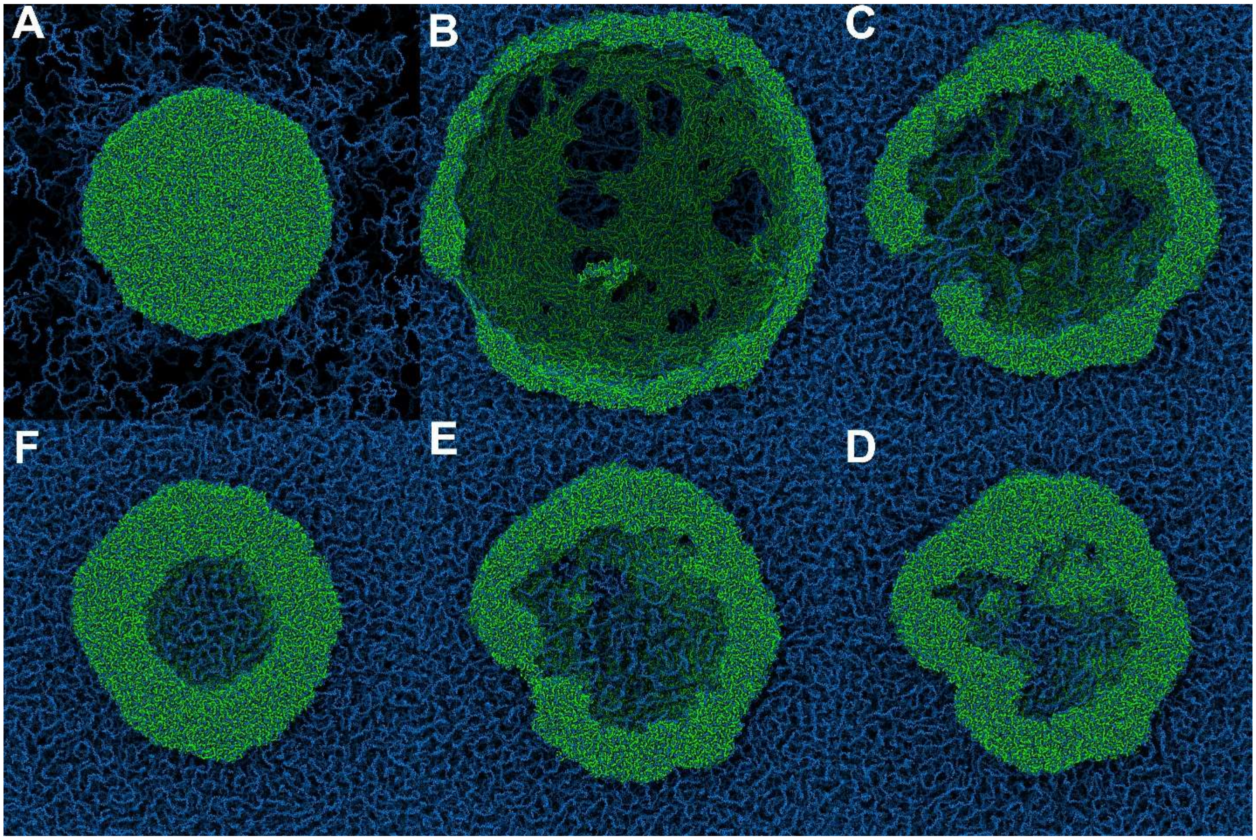


**Fig. S15.** MD simulation snapshots showing the transition from micelle to vesicular structure. We consider a micelle formed and equilibrated at poly(U) RNA concentration of 0.70 mg/ml. The RNA concentration in the simulation box is increased to 14.60 mg/ml by adding RNA around the micelle. The system is equilibrated in NVT ensemble at 298 K and the snapshots are taken at (***A***) 0.0, (***B***) 0.3, (***C***) 0.6, (***D***) 0.9, (***E***) 1.2 and (***F***) $10 \times 10^5 \tau$ time steps (also see Movie S7).

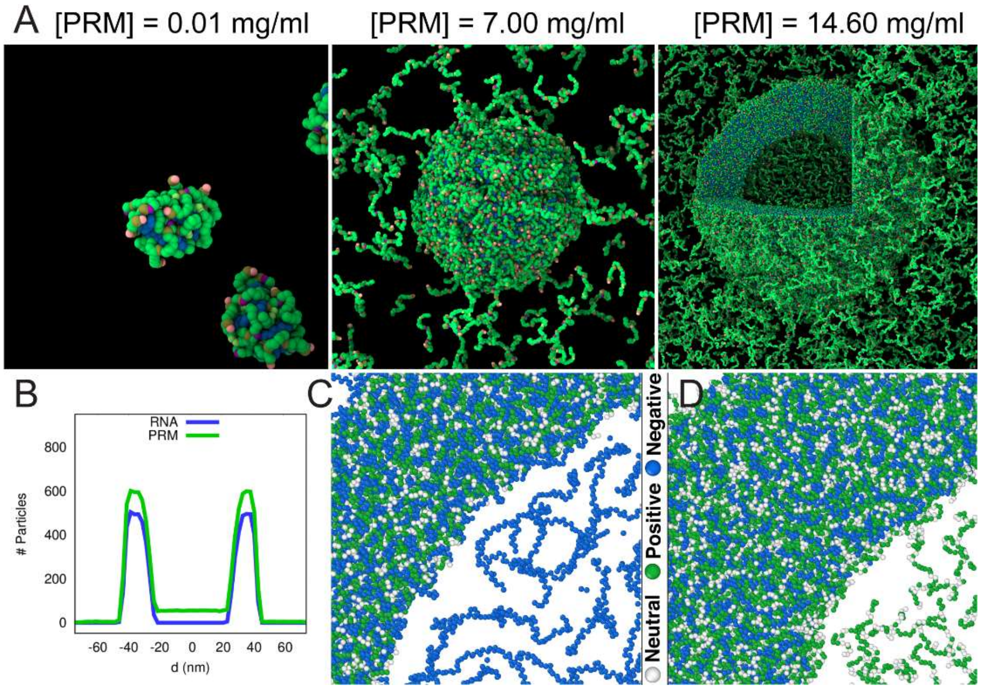


**Fig. S16.** ***(A)*** Equilibrium MD configurations showing tadpole, micelle, and vesicle structures formed under charge disproportionate PRM excess conditions, $Conc_{PRM} = 5 \times Conc_{RNA}$. PRM: *green*; and RNA: *blue*. See also Figure 3*B* in the *Main-text*. The concentration of PRM used are indicated in each snapshot. (***B***) Density profiles of PRM and RNA for the vesicle structure formed in excess PRM conditions as computed from the MD trajectory. Close up views of vesicle rims in ***(C)*** excess RNA, and ***(D)*** excess PRM conditions.

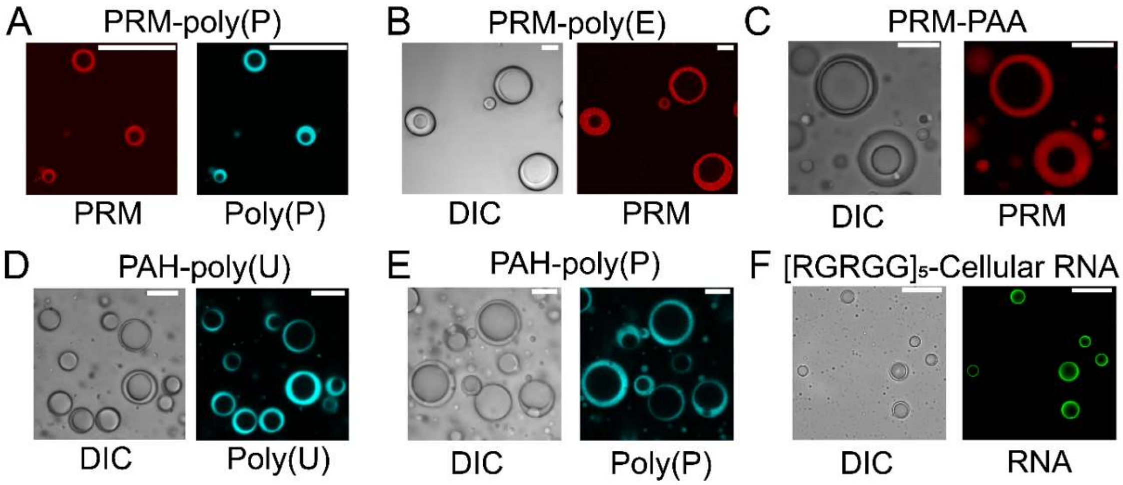


**Fig. S17.** Bright-field and fluorescence images of vesicles formed by ***(A)*** Protamine (PRM) and polyphosphate [poly(P)] mixed at concentrations of 4.4 mg/ml and 22 mg/ml, respectively. (Buffer: 15 mM Tris-HCl, pH 7.5). ***(B)*** Protamine (PRM) and poly(Glutamic acid) [poly(E)] mixed at concentrations of 17.6 mg/ml and 18 mg/ml, respectively. (Buffer: 10 mM Tris-HCl, pH 7.5). ***(C)*** Protamine (PRM) and poly(acrylic acid) (PAA) mixed at concentrations of 8.8 mg/ml and 22 mg/ml, respectively. (Buffer: 15 mM Tris-HCl, pH 7.5). ***(D)*** poly(Allylamine) (PAH) and poly(U) RNA at concentrations of 40 mg/ml and 4 mg/ml, respectively. (Buffer: 25 mM Tris-HCl, pH 7.5). ***(E)*** poly(Allylamine) (PAH) and polyphosphate (polyP) mixed at concentrations of 70 mg/ml and 4 mg/ml, respectively. (Buffer: 5 mM Tris-HCl, pH 7.5). ***(F)*** $[RGRGG]_5$ and yeast total RNA (cellular RNA) mixed at concentrations of 0.024 mg/ml and 8.9 mg/ml, respectively. (Buffer: 5 mM Tris-HCl, pH 7.5). Probes used are PRM-Alexa594 for PRM-containing vesicles and DAPI for poly(U) RNA and polyphosphate-containing vesicles. SYTO13 was used to image vesicles formed by cellular RNA. Scale bars represent 20 μm.

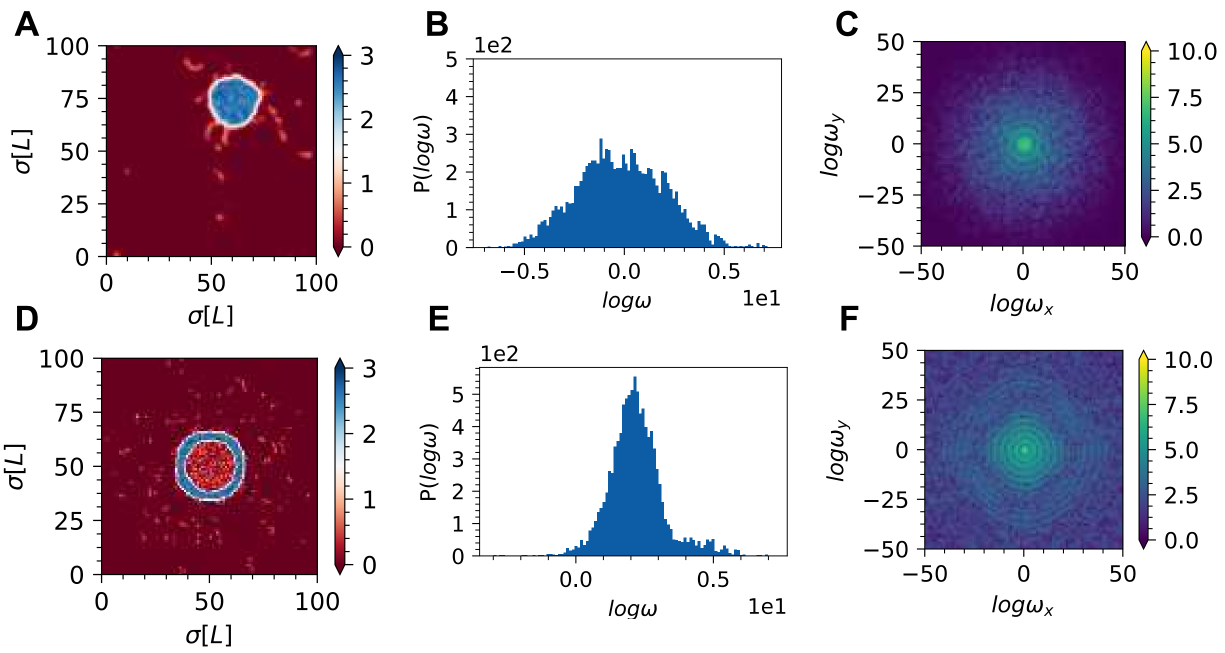


**Fig. S18. Power spectral analysis of RNA ordering in (A-C) micelles and (D-F) vesicles.** (***A*** and ***D***) Gaussian density of ensemble averaged RNA positions calculated for micelle and vesicle states respectively. (***B*** and ***E***) 1D Power spectrum of RNA position density calculated via Gaussian blurring of Cartesian positions of monomers in RNA chains calculated for micelle and vesicle states respectively. (***C*** and ***F***) 2D power spectrum calculated for micelle and vesicle states respectively.

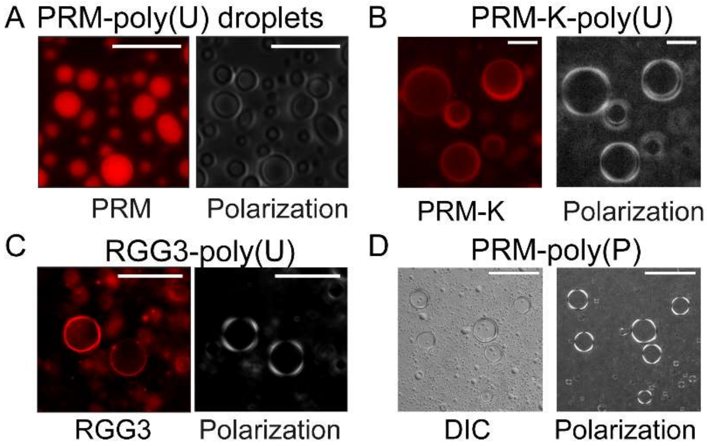


**Fig. S19.** Cross-polarization light microscopy images of ***(A)*** PRM and poly(U) droplets [0.44 mg/ml PRM and 0.22 mg/ml poly(U)], ***(B)*** PRM-K-poly(U) vesicles [4.4 mg/ml PRM-K and 22 mg/ml poly(U)], ***(C)*** RGG3-poly(U) vesicles [4.0 mg/ml RGG3 and 20 mg/ml poly(U)], and ***(D)*** PRM-polyP vesicles (4.4 mg/ml PRM and 22 mg/ml polyP). All samples were prepared in 15 mM Tris-HCl buffer (pH 7.5). Scale bars represent 10 μm for ***A***-***C*** and 100 μm for ***D***.

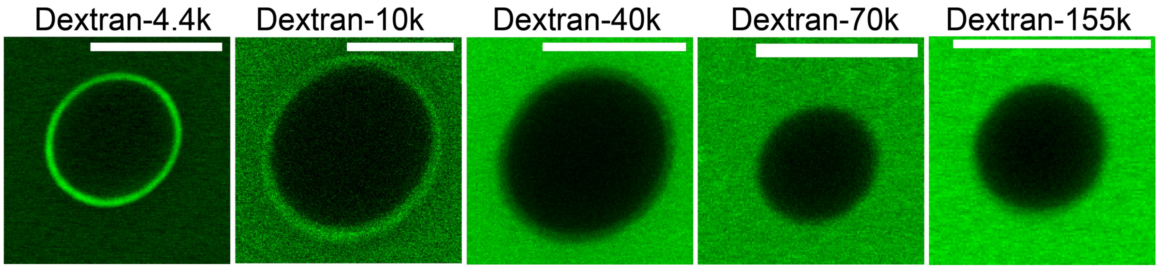


**Fig. S20.** Dextran partitioning in PRM-poly(U) vesicle rim and lumen as a function of dextran molecular weight. Samples are made at 4.4 mg/ml and 22 mg/ml poly(U) in 15 mM Tris-HCl buffer. Scale bars represent 10 μm. The corresponding summary plot is shown in Figure 5*C* in the *Main-text.*

### Supplementary Movie Legends

**Movie S1 (separate file).** FRAP movie of PRM-poly(U) hollow condensate membrane. Scale bar represents 10 µm.

**Movie S2 (separate file).** Optical tweezer-controlled fusion of two PRM-poly(U) vesicles. Scale bar represents 10 µm.

**Movie S3 (separate file).** Optical tweezer-controlled fusion of a PRM-poly(U) droplet with a PRM-poly(U) hollow condensate. Scale bar represents 10 µm.

**Movie S4 (separate file).** Droplet to vesicle transition induced by flowing RNA at high concentration into a microfluidics channel. Scale bar represents 10 µm.

**Movie S5 (separate file).** Hollow condensate to droplet transition induced upon RNase-A addition. Scale bar represents 10 µm.

**Movie S6 (separate file).** MD trajectory showing formation of micelles from tadpoles during an NVT simulation at 298 K.

**Movie S7 (separate file).** MD trajectory showing formation of vesicle from micelle during an NVT simulation at 298 K. We consider a micelle formed and equilibrated at RNA concentration of 0.70 mg/ml. The RNA concentration in the simulation box is increased to 14.60 mg/ml by adding RNA around the micelle. The movie shows the spontaneous formation of a vesicle in response to the increase in external RNA concentration.

### Note-1

Our experimental results (Fig. 2A in the *Main-text*) can be summarized by the following two key observations on the location of hollow condensates within the state diagram: (i) protein-RNA complexes form micellar and vesicular condensates at disproportionate mixture compositions, (ii) at intermediate volume fractions, the micellar phase is stabilized whereas above a threshold concentration, the vesicular phase is stabilized. In parallel, our molecular dynamics simulations reveal similar observations with the addition of tadpole-like structures appearing at relatively lower volume fractions. These data collectively suggest a qualitative state diagram as shown in Figure S21. We note that tadpole-like structures at disproportionate mixture compositions were originally predicted by Zhang and Shklovskii using a phenomenological theory of polyelectrolyte condensation (10). Furthermore, tadpole-like structures were experimentally observed for DNA-Chitosan systems at non-stoichiometric mixing conditions (11). Here, we propose a generic phase behavior for *mixtures of polymers that undergo phase separation via obligate heterotypic interactions: at disproportionate mixture compositions, they form anisotropic tadpole-like complexes at low chain concentrations, micellar condensates at intermediate chain concentrations, and vesicles at high chain concentrations.*

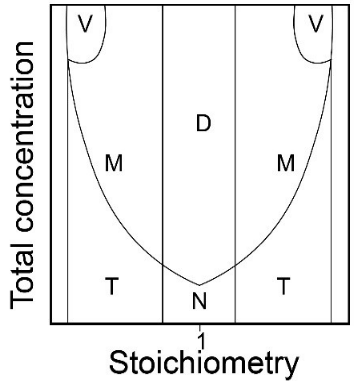


**Fig. S21.** A qualitative phase diagram summarizing our observations. **T**: tadpoles, **N**: neutral and quasi neutral linear polyelectrolyte complexes, **M**: spherical micelle regime, **D**: neutral droplets formed by phase separation, **V**: Vesicle phase.

The existence of a structural transition in the system with increasing concentrations suggests a major role of non-monotonic variations in interactions of heterotypic complexes with each other as well as with the solvent. The configuration and arrangement of chains in a given phase at equilibrium is the one that has lowest free energy. In this note, we qualitatively rationalize the state diagram shown in Figure S21 in light of the current theories of heterotypic condensation. We start by analyzing the tadpole formation at disproportionate mixture compositions as a result of differential solvation. Next, we describe the tadpole-to-micelle transition by drawing analogies with scaling and phenomenological theories. Similar analysis is also presented for the micellar to vesicular transition. Finally, we summarize and discuss inadequacies in our current understanding of the phenomenon of hollow condensate formation. While we realize that the mechanism and driving forces for vesicle formation warrants a separate and deeper theoretical/computational investigation, *the evidence presented in this work suggests that hollow condensates are a manifestation of non-stoichiometric anisotropic complexes with non-uniform solvation properties. These complexes can form via heterotypic interactions between associative polymers at far from stoichiometric mixing ratios* (12, 13).

#### Low concentration regime: Formation of tadpoles

Consider mixture of associative polymers with stickers-and-spacers architecture (12-14), such as a positively charged RNA-binding protein and a negatively charged RNA at disproportionate mixture composition. We consider a super-stoichiometric condition where the concentration of RNA is much higher than the concentration of the protein. For each RNA chain, there is a limited number of proteins such that each RNA chain can only be partially occupied by proteins. Shklovskii and Zhang argued that in such cases, the proteins may only preferentially screen one side of the RNA chain (Fig. S22*A*). This preferential screening at the open ends has been argued for weakly charged polyelectrolytes using scaling models and for strongly charged polyelectrolytes using molecular dynamics simulations (15, 16). One of the consequences of such preferential screening of the chain is the formation of tadpole-like structures where a partial nano-condensate is formed as a result of short-range attractions due to the bound protein molecules (10). Consequently, the tadpole structure can be described as having a neutral head and a charged tail (Fig. S22*B*). Independent of Shklovskii-Zhang model, the formation of these “tadpoles” can also be rationalized by multiple other theoretical frameworks. For example, Shusharina and Rubinstein used scaling theories to show the formation of tadpoles by di-block copolymers containing a neutral block and a charged block (17). In light of the preferential screening idea, one can approximate a partially complexed RNA chain as a di-block copolymer (Fig. S23). Following Shusharina and Rubinstein, the partially complexed RNA chains will form tadpoles to reduce the interface between the neutralized block and the solvent (Fig. S22*C*). Here, we hypothesize that the neutralized segment (block B; Fig. S23) has less-favorable interactions with the solvent than the bare segment (block A; Fig. S23). Similar results can be derived from calculating the effective solvation volume following Harmon, Holehouse, and Pappu (13). Briefly, Harmon *et al.* showed that having polymers with different effective solvation volume interacting with a common partner can lead to the formation of multi-layered condensates, where the less soluble polymer forms the core and the more soluble polymer forms the shell of the condensate. The authors used this idea to argue that multilayered condensates can be formed by changing the solvation volume of a fraction of the protein population through post-translational modifications such as phosphorylation. Phosphorylation at a given site increases charge on the protein chains and may result in increasing its solubility (*i.e.*, higher solvation volume compared to a non-phosphorylated chain). Therefore, phosphorylation of a fractional population of proteins can lead to the formation of multi-layered condensate with the phosphorylated chains residing at the outer shell of the condensate and non-phosphorylated chains at the inner core (in presence of a common multivalent partner). In a similar vein, we can argue that having a partially complexed RNA chain means that the effective solvation volume of the bare segment is larger than that of the protein-bound segment of the chain. This scenario may explain the formation of a “tadpole head” by the protein-bound part and “tadpole tail” by the bare segment of the chain (Fig. S22). An important distinction between the system studied by Harmon *et al.* and our system is that the difference in solvation volume in their case is from chain to chain (*i.e.*, inter-chain differential solvation, such as a phosphorylated chain and an un-phosphorylated chain) while the difference in our case is from distinct segments within the same chain (intra-chain differential solvation).

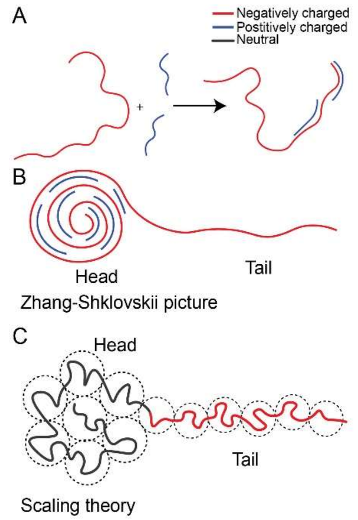


**Fig. S22.** ***(A)*** Partially screened complex. ***(B)*** A tadpole in Zhang-Shklovskii picture (strong polyelectrolytes), and ***(C)*** the scaling picture of Susharina and Rubinstein (weak polyelectrolytes).

The aforementioned arguments suggest that tadpole formation can be rationalized by an intra-chain differential solvation due to preferential screening of a part of the RNA chain by the bound proteins. This preferential screening could be a generic result of protein-RNA obligate heterotypic interactions. The fact that these structures can be rationalized by theories built on distinct sticker properties (such as weak *vs.* strong polyelectrolytes) implicates that they maybe generic and resilient over a wide range of solution conditions (such as buffer ionic strength). In what follows, we argue that tadpoles can self-assemble into spherical micelles and hollow condensates, the latter was observed to be persistent over a wide range of salt concentrations (25 to 500 mM NaCl; Fig. S2).

**Intermediate concentration regime: Formation of micelles**

Our MD simulation suggests that the micellar condensates are formed by coalescence of tadpoles at intermediate volume fractions of the protein and the RNA chains (see Figure 3 in *Main-text* and Movie S6). This is corroborated by the observation of small spherical condensates in our experiments (see Fig. 1*B*, *Main-text* and Fig. S12). To rationalize micelle formation, we hypothesize the following: increasing the total volume fraction of the RNA-protein complexes at disproportionate mixture composition leads to an increase in the concentration of tadpoles. Above a critical concentration, tadpoles form spherical micelles with a core and a corona. The core is enriched in tadpoles' heads and the corona is formed by the tadpoles' bare tails. The driving force of micelle formation is the reduction in the interfacial contact between the tadpole heads and the solvent upon forming the micelle. The opposing force for this condensation of heads is the repulsive energy between tadpole tails as well as the mixing entropy. A critical micelle concentration ($cmc$) can be defined as the concentration of tadpoles at which these two forces balance each other. Shusharina and Rubinstein derived the $cmc$ for their $neutral-charged$ di-block copolymers as follows (17):

$$cmc\, b^3 = \exp\left(-\frac{\Delta F}{k_B T}\right) = \exp\left(-\left(1 - p^{-\frac{1}{3}}\right) N_B^{\frac{2}{3}} |v|^{\frac{4}{3}}\right) \quad (1)$$

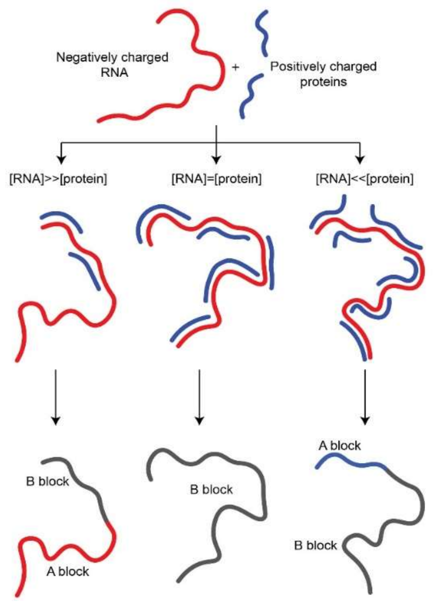


**Fig. S23.** Complexation of polyelectrolyte at various charge disproportionation states. Preferential screening of the chain end leads to a complex that can be approximated as a di-block copolymer with a charged A block and a neutral B block. (red) negatively charged segments. (blue) positively charged segments.

Where $p$ is the number of tadpoles in one micelle, $N_B$ is the length of the neutral block and $v$ is the excluded volume parameter. Note that under the approximation we made earlier (Fig. S23), the size of the tadpole head increases as the ratio between protein and RNA approaches the stoichiometric ratio. This means that the closer

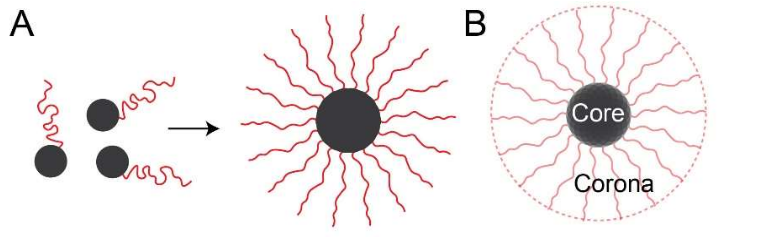


**Fig. S24.** ***(A)*** Tadpoles condense into spherical micelle with brush-like architecture. ***(B)*** The micelle is composed from the neutral core and the charged corona.

the system is to stoichiometric mixture composition, the larger is the relative size of the neutral block (tadpole head in our case) and hence, the lower is the $cmc$ [see equation (1)].

In summary, we argue that tadpoles form spherical micelles above a threshold concentration due to the reduction of the interface between the tadpole's head and the solvent upon micelle formation. The resulting micelles have a core that is enriched in the protein-bound segments of the chain and a corona that consists of the bare part of the chain. This structure is qualitatively similar to what *Harmon et al.* predicted, *i.e.*, the more soluble charged segments will form the shell and the less soluble neutral segments will form the core (13) (Fig. S24).

**High concentration regime: Formation of vesicles**

Our experimental state diagram indicates that vesicle formation occurs at compositionally disproportionate mixtures with a high total volume fraction of protein and RNA chains. This result is also supported by our molecular dynamic simulations (see Fig. 3, *Main-text*). To understand vesicle formation in light of the current model, we consider two experimentally adjustable parameters: mixture composition and the total volume fraction of the protein-RNA complexes. In the state diagram presented in Figure S25, these two parameters are plotted along the X- and Y-axis, respectively. Let us consider a micelle that is formed at a specific RNA-to-protein ratio under RNA-excess condition. Adding additional RNA to the external dilute phase will create a competition between RNA chains for binding to fewer available proteins per RNA. This process would result in the redistribution of protein chains along an RNA chain in a way that will result in tadpoles with shorter heads and longer tails on average. This means that the repulsion on the micellar surface which opposes the formation of a micelle will increase, and thereby destabilizing the micellar phase (S26*A*, (ii)). Under this condition: two scenarios are possible: (a) the micelle will disassociate into a solution of tadpoles, and/or, (b) the micelles will undergo a structural transition that will result in lowering the overall free energy of the system (Fig. S26*A*, (i) and (iii)). Our experiments and simulations clearly suggest that under such a condition, the micellar condensates transition to a vesicle-like structure. These results indirectly indicate that the disassociation of micelles into tadpoles will increase the free energy of the system, which could well be through increasing the interfacial interactions of the tadpole heads with the solvent. On the contrary, the hollow condensates provide an effective way to reduce their solvation volume as well as interactions between tadpole tails (Fig. S26*A*, (iii)) by providing a larger surface area (through the creation of a secondary surface). The described picture is corroborated in MD simulations showing explosive transition of a single micelle to a vesicle upon embedding the micelle in a large bath of RNAs (Movie S7; Fig. S15). Similarly, our microfluidics experiment where we optically trap a droplet and flow excess RNA also revealed a similar transition (Fig. S11).

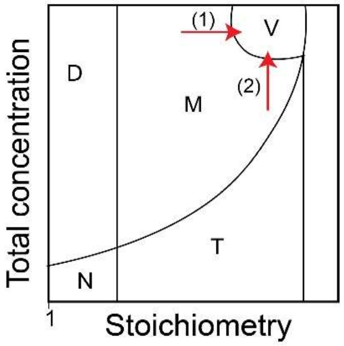


**Fig. S25.** Two pathways to reach the vesicle phase in the phase diagram: (1) Increasing charge disproportionation and (2) Increasing total concentration of proteins and RNA.

The second parameter that plays an important role in vesicle formation is the total concentration of protein and RNA chains (*i.e.*, adding more tadpoles, Figs. S25 & S26*B*). Increasing the total concentration of proteins and RNAs without changing the mixture composition would lead to an increase in the number density of micelles. We hypothesize that increasing the number density of protein-RNA micelles mediates a structural transition of micelles into vesicles. This is verified by our experiments and simulations (Figs. 2*A* & 3, *Main-text*). We suggest that an additional interface in the vesicles is the driving force underlying this structural transition. The creation of an additional interface provides larger solvation volume for the bare RNA chains without increasing the exposure of the neutralized segments to the solvent (Fig. S26*B*, (iii)). Without such a structural transition, the increased overlap between micelle coronas at higher micellar density will result in a decrease in the solvation of bare RNA chains and will destabilize the system by increasing its total free energy.

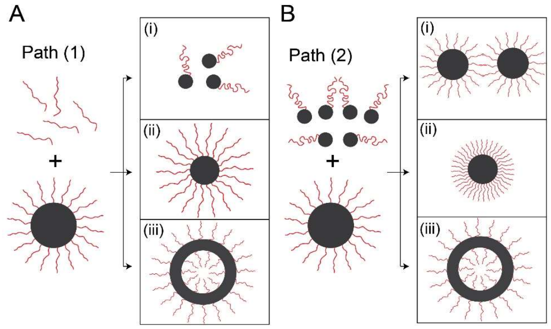


**Fig. S26.** Possible scenarios for the two pathways highlighted in Figure S25. In both these pathways, the hollow condensate provides the configuration with the lowest free energy (see text).

### Concluding remarks

We note that the proposed mechanism to rationalize our observations from experiments and simulation is strictly phenomenological. Moreover, the model considered here emphasizes the asymmetry in the length of the associative polymers. However, our experiments revealed vesicle formation for ternary mixtures with components that have lesser length asymmetry (see Fig. 4*C*, *Main-text*) and systems with large polydispersity such as $(RGRGG)_5$-cellular RNA mixtures (Fig. 4*F*, *Main-text*). We speculate that such systems may form different kinds of micellar structures than simple tadpoles, including dimers or star-like complexes at disproportionate mixture compositions (18). The tadpole picture seems to be consistent with reentrant phase transitions, but may be counterintuitive for mixtures with equal length polymers or when the component in excess is the shorter polymer. The formation of vesicles by a number of ternary systems where phase separation is governed by heterotypic interactions (Fig. 4, *Main-text*) points out that this is a general phenomenon in obligate heterotypic complexation. Recent theoretical work already shows that such systems undergo reentrant liquid condensation (12), implying that hollow condensate formation may be common in a vast array of biological and synthetic systems.
